# Molecular Basis of the Activation and Dissociation of Dimeric PYL2 Receptor in Abscisic Acid Signaling

**DOI:** 10.1101/721761

**Authors:** Chuankai Zhao, Diwakar Shukla

## Abstract

Phytohormone abscisic acid (ABA) is essential for plant responses to biotic and abiotic stresses. Dimeric receptors are a class of ABA receptors that are important for various ABA responses. While extensive experimental and computational studies have investigated these receptors, it remains not fully understood how ABA leads to their activation and dissociation for interaction with downstream phosphatase. Here, we study the activation and the homodimeric association processes of PYL2 receptor as well as its heterodimeric association with the phosphatase HAB1 using molecular dynamics simulations. Free energy landscapes from ~223 *μ*s simulations show that dimerization substantially constrains PYL2 conformational plasticity and stabilizes inactive state, resulting in lower ABA affinity. Also, we establish the thermodynamic model for competitive binding between homodimeric PYL2 association and heterodimeric PYL2-HAB1 association in the absence and presence of ABA. Our results suggest that the binding of ABA destabilizes PYL2 complex and further stabilizes PYL2-HAB1 association, thereby promoting PYL2 dissociation. Overall, this study explains several key aspects on activation of dimeric ABA receptors, which provide new avenues for selective regulation of these receptors.

## Introduction

Abscisic acid (ABA, Fig. 1A) is a vital plant hormone that responds to a variety of environmental stresses, including drought which significantly impacts crop yield worldwide.^1,2^ When plants are under water deficiency, *in planta* biosynthesis of ABA is promoted^3^ and elevated levels of ABA eventually lead to stomata closure, thereby reducing transpiration water loss.^4^ ABA responses are controlled by a negative regulatory signaling network that involves soluble PYR/PYL/RCAR (pyrabactin resistance1/PYR1-like/regulatory component of ABA receptor) ABA receptors, PP2Cs (clade A serine/threonine protein phosphatases 2C), and SnRK2s (subfamily 3 SNF1-related kinase 2).^5^ In the absence of ABA, PP2Cs dephosphorylate and inactivate SnRK2s (Fig. 1B).^6,7^

**Fig. 1.**
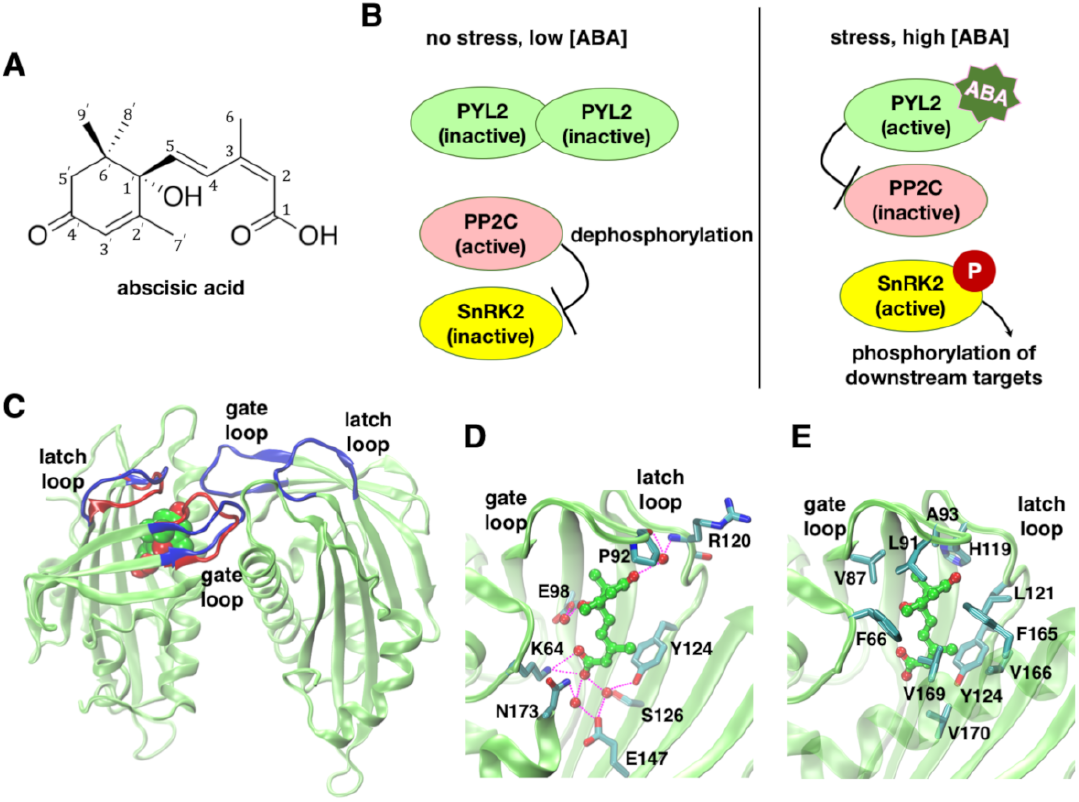
ABA signaling mechanism and PYL2 receptor structure. (A) ABA molecule and (B) the core components involved in ABA signal transduction. (C) ABA binds to PYL2 receptor and triggers the conformational changes in the gate loop and the latch loop. The inactive and active (PDB ID: 3KDH and 3KDI^29^) loop conformations are shown in blue and red, respectively. (D) Polar and (E) van der Waals interactions between ABA and the amino acids in PYL2.

When ABA binds to PYR/PYL/RCAR receptors, these receptors inactivate PP2Cs via stable binding and prevent SnRK2s from dephosphorylation by PP2Cs.^8^ SnRK2s then activate through autophosphorylation, which triggers the downstream signaling cascade (Fig. 1B).^9–11^ PYR/PYL/RCAR receptors are promising targets for improving plant water use efficiency.^12–15^ Notably, a series of ABA agonists covering various chemotypes^8,16–21^ and a variety of engineered ABA receptors^22,23^ have been discovered in recent years, in order to achieve selective activity control of PYR/PYL/RCAR receptors in multiple plant species.

In *Arabidopsis thaliana*, PYR/PYL/RCAR receptors can be classified according to their oligomeric states in solution: PYR1, PYL1 and PYL2 form a stable homodimer, while PYL4-12 are monomeric and PYL3 is in equilibrium between monomeric and dimeric states.^24–27^ PYL1-13 share 50-73% sequence similarity and 38-64% sequence identity as compared to PYR1 and these receptors have identical fold (Fig. S1). Since 2009, the gatelatch-lock mechanism for explaining the mode of action of ABA has been established from crystallographic structural studies on ABA receptors (Fig. 1C).^24,25,28,29^ ABA binds to a structurally conserved hydrophobic binding pocket which is filled by water molecules.^24^ Upon ABA binding, two flexible loops at top of the binding cavity, named as the gate loop and the latch loop, will undergo large conformational changes to close and activate ABA receptors (Fig. 1C).^28^ In PYL2, the binding of ABA is stabilized by both direct and water-mediated hydrogen bonding interactions with the polar residues (K64, Y124, S128, E147, N173) at bottom of the binding site (Fig. 1D) as well as hydrophobic interaction with the non-polar residues (Fig. 1E).^24^ The closed gate loop creates interaction interface to facilitate the binding between ABA receptors and PP2Cs. The monomeric receptors show ABA-independent constitutive activity,^26,30^ whereas the dimeric receptors are ABA-dependent and have nearly two orders of magnitude lower ABA affinity.^31^ When PP2Cs bind to ABA-bound receptors, PP2Cs form water-mediated interaction with the carbonyl group of ABA. ^28^ Therefore, PP2Cs are often considered as ABA co-receptors. ^32^

While a large number of experimental studies have probed the ABA signaling mechanism, the atomic-scale dynamics of ABA perception and subsequent conformational changes in ABA receptors remained elusive. Specifically, it was unclear how the binding of ABA happens and leads to the activation of PYR/PYL/RCAR receptors, and how these processes vary among the receptors. In our recent work^33^, we have employed molecular dynamics (MD) simulations to investigate ABA-mediated activation processes of monomeric PYL5 and PYL10 belonging to two separate clades of PYR/PYL/RCAR receptors in *Arabidopsis thaliana*. Strikingly, the gate loop remains flexible between open and closed conformations after ABA binding, in agreement with the hydrogen/deuterium exchange mass spectrometry experiment on ABA receptor dynamics and the intermediate ABA-bound receptor crystal structure with the gate loop open.^34,35^ ABA binding is thus necessary but insufficient for full activation of monomeric receptors. PP2C binding stabilizes the closed gate loop conformation, which explains enhanced ABA affinity in the presence of PP2C. ^36^ In contrast to PYL5, the gate loop in PYL10 can close without ABA binding, which unravels the PYL10 ABA-independent activation mechanism.^26,30^

Previous studies have shown that dimeric receptors are critical for ABA responses, despite their lower ABA sensitivity compared to monomeric receptors. For example, activation of dimeric receptors by selective ABA agonists has led to guard cell closure in both soybean and *Arabidopsis thaliana*.^16,18^ Several questions remain unanswered regarding the activation mechanism of dimeric ABA receptors. Firstly, it remains not fully understood how the dimerization of ABA receptors leads to ABA-dependent receptor activation and lower ABA affinity in contrast to their monomeric counterparts. While dimeric receptors such as PYL2 are inactive in the absence of ABA, it was shown that monomeric variant of PYL2-I88K obtained weak constitutive activity on PP2C inhibition. ^26^ Furthermore, ABA binding affinity to PYL2-I88K is 7-fold higher compared to wild-type PYL2.^26^ What is the molecular origin of intrinsic ABA affinity differences between monomeric and dimeric receptors? How does ABA bind to dimeric receptors and mediate their conformational changes? Secondly, it is unclear how ABA affects the assembly of dimeric receptors and the heterodimeric association between these receptors and PP2Cs. To trigger the downstream signaling, ABA receptor is required to inhibit PP2C by forming a 1:1 heterodimer. However, the homodimer interface of dimeric receptors and the receptor-PP2C heterodimer interface overlap with each other, which excludes the possibility of the occurrence of a trimeric complex. Dimeric receptors thus need to dissociate prior to heterodimerization with PP2C. However, PYR1 and PYL1-2 remain in a dimer conformation in the presence of ABA suggested by a range of experimental characterizations.^25,27,29,31^ It was noted that the gate loop closure caused by ABA binding may weaken dimeric receptors by comparing the structures of ABA-free and ABA-bound PYL2 crystal structures.^29^ Overall, the molecular mechanism of activation and dissociation of dimeric ABA receptors requires further investigation.

In this work, we seek to understand the molecular mechanism of activation and dissociation of PYL2 receptor using various computational approaches (Table S1). We have utilized extensive all-atom MD simulations (~223 *μ*s aggregate) to elucidate the molecular basis of ABA-mediated PYL2 activation in both its monomeric and dimeric forms. Using Markov state models (MSMs) to analyze massively parallel MD trajectories,^37–41^ we have elucidated the full pathway of ABA binding and PYL2 activation, along with quantitative thermodynamic and kinetic characterizations. To understand the role of ABA in homodimeric PYL2 association and heterodimeric PYL2-PP2C association, we have utilized replica exchange umbrella sampling (REUS) simulations^42^ and multistate Bennett acceptance ratio (MBAR) method^43^ to accurately estimate the standard association free energies for both cases.^44–46^ We have established a thermodynamic model to explain the competitive interactions between PYL2 self-association and its association with the downstream phosphatase HAB1. Overall, this study provides key molecular insights into the activation of an important class of ABA receptors, which can be exploited for future genetic and agrochemical control of ABA-regulated stress responses.

## Results and discussion

### Simulations reveal the conformational landscapes of the monomeric and the dimeric PYL2

We have performed ∼ 107 *μ*s and ~116 *μ*s (aggregate) adaptive ensemble MD simulations^47–52^ to study ABA binding and subsequent conformational changes in both the monomeric and the dimeric PYL2, respectively (Tables S2 and S3). While PYL2 only exists as a dimer in solution, we study the dynamics of both the monomeric and the dimeric states in order to investigate how the dimerization affects PYL2 activation. The simulations are initialized from the inactive PYL2 crystal structure (PDB: 3KDH^29^). In both cases, a single ABA molecule is randomly placed far away from the binding site. We perform multiple rounds of short parallel MD simulations to achieve ABA binding and subsequent conformational changes in PYL2. For the dimeric PYL2, our simulations have captured the binding of ABA to one of the protomers in PYL2 and the closure of the gate loop in the ABA-bound protomer (Fig. S2A). In principle, ABA can bind to either protomer without any preference due to the symmetry of the homodimer structure. We have focused our analysis on the dynamics of the ABA-bound protomer as captured in our simulations. We use Markov state models (MSMs) to analyze the sampled PYL2-ABA conformational space via long timescale MD simulations. MSMs allow to aggregate all individual trajectories and represent the full conformational space as discretized microstates and transitional probabilities between these states.^37–39^ All the conformations collected from MD simulations are clustered into 300 and 200 discretized states for the monomeric and the dimeric PYL2, respectively. The features for clustering PYL2-ABA conformations are 32 distances for the monomeric PYL2 and 64 distances for the dimeric PYL2, which are used to describe the conformations of both the gate loop and the latch loop, ABA position and PYL2-ABA interactions (Table S4 and Fig. S2B). MSMs are then built to estimate the equilibrium probabilities of all states and their inter-state transition probabilities. The thermodynamics and kinetics of ABA binding and receptor activation processes can be inferred from these MSMs.^38,39^ Details of the simulations and analysis are summarized in Supplementary Methods.

We report the two-dimensional conformational free energy landscapes for both the monomeric and the dimeric PYL2 receptors (Fig. 2A, B) as well as the associated error bars on the landscapes (less than 0.4 kcal/mol, Fig. S3A, B). The free energy landscape is a weighted projection of the sampled protein-ligand conformations onto relevant structural metrics, which show the equilibrium probability of the conformational states. By inspecting the free energy landscape, one can identify the important protein-ligand conformation states and investigate the thermodynamic process associated with the transition between these states. The x and y axes of the landscapes are the distance between PYL2 and ABA and the root mean square deviation (RMSD) of the gate loop from active PYL2 structure, which indicate whether ABA is bound to PYL2 and whether the gate loop is in open or closed states, respectively. The free energy landscapes of both receptors share similar minima, which correspond to the binding intermediate states and the ABA-bound states. Before ABA binds, there is only a single flattened minima on both the landscapes, with the gate loop RMSD centering around ~4-5 Å indicating an open gate loop conformation. The free energies of these binding intermediate states are 0-2 kcal/mol, suggesting that they are relatively stable conformations. However, the gate loop RMSD from the active structure fluctuates between 1-11 Å for the monomeric PYL2, whereas the gate loop RMSD for the dimeric PYL2 only fluctuates between 2-8 Å. These results show that breaking the PYL2 complex results in a higher degree of conformational flexibility of the gate loop. Since the gate loop in monomeric PYL2 may adopt a closed conformation before ABA binds, the monomeric variant of PYL2 receptor would exhibit weak constitutive activity in solution. Our results are in agreement with the finding that the monomeric PYL2-I88K exhibited an increased constitutive activity on PP2C inhibition compared to the dimeric PYL2.^26^

**Fig. 2.**
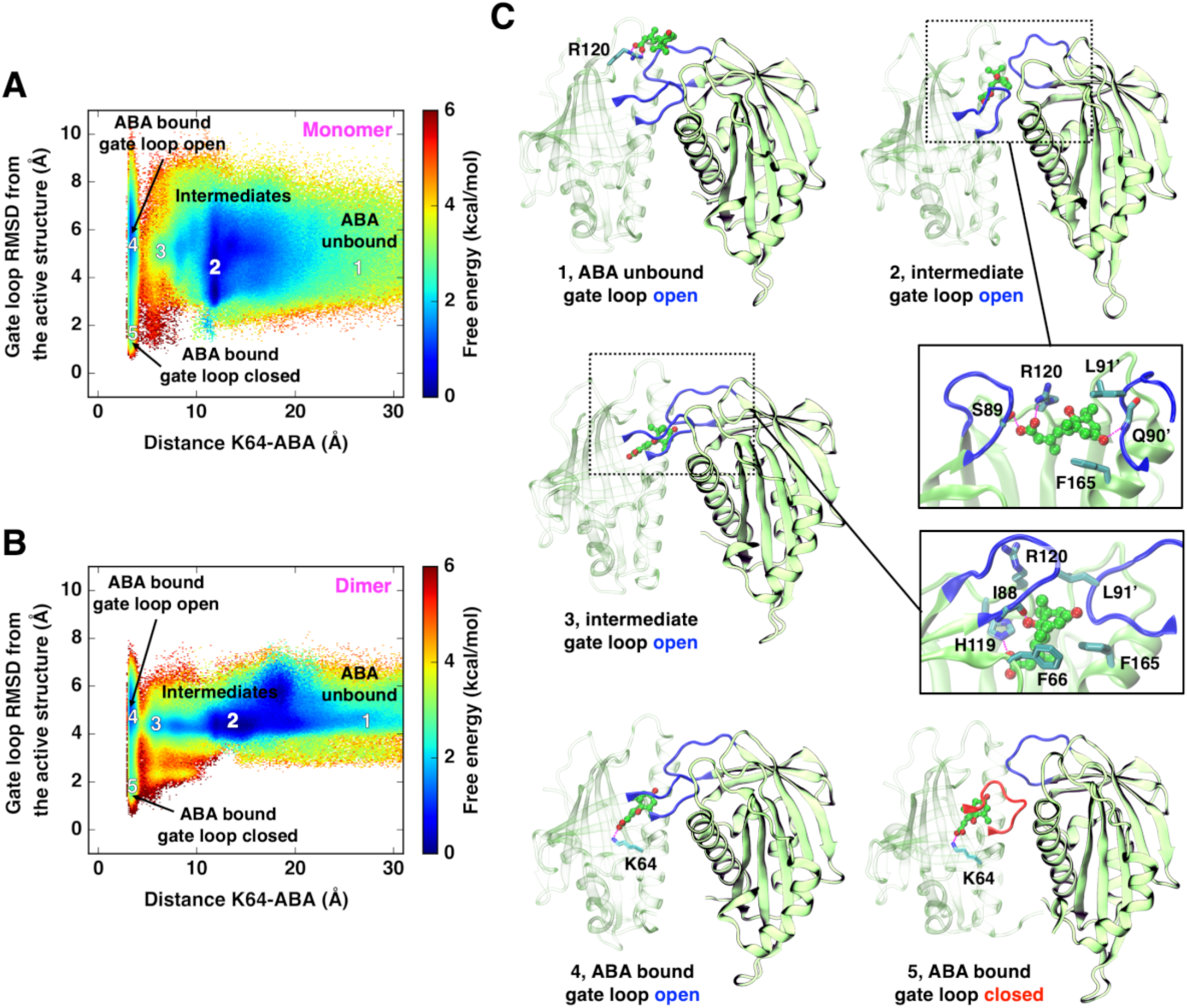
ABA binding and subsequent PYL2 activation processes. Free energy landscapes of ABA binding to (A) the monomeric and (B) the dimeric PYL2 receptors. The landscapes are generated by projecting all the conformations onto two metrics, the distance between ABA and the binding site (K64 in PYL2) as well as the gate loop RMSD from the PYL2 active crystal structure (PDB: 3KDI^29^), weighted by MSM probabilities. K64-ABA distance is measured as distance between the carbon atom in -COO^−^ group of ABA and the NZ atom in 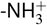 group of K64. The *C_α_* atoms of V87-S92 in PYL2 are used for calculating RMSD of the gate loop. (C) Snapshots of the transition states 1-5 in the pathway of ABA binding and PYL2 activation as identified via transition path theory analysis. The states 1-5 are mapped onto (A) and (B) based on their corresponding K64-ABA distances and gate loop RMSDs. Key residues that interact with ABA are shown, and the gate loops are highlighted in blue (open) or red (closed).

After ABA binds, we observe two minima that correspond to the open and closed gate loop conformations (Fig. 2A, B). The landscapes clearly suggest that the binding of ABA promotes the gate loop closure. For the monomeric PYL2, the free energies for the open gate loop and the closed gate loop states are nearly comparable (<2 kcal/mol), suggesting that the gate loop is in equilibrium between the open and the closed conformations after ABA binding. This is consistent with our previous finding on the monomeric PYL5 and PYL10 that ABA binding is necessary but insufficient for full receptor activation. ^33^ In contrast to the monomeric PYL2, the free energy of the open gate loop state for the dimeric PYL2 (<2 kcal/mol) is lower as compared to that of the closed gate loop state (~3 kcal/mol). In other words, the open gate loop conformation is stabilized due to the formation of dimer structure. Our results are further supported by the published hydrogen/deuterium exchange (HDX) mass spectrometry data on the PYL2 gate loop dynamics in solution. ^34^ West *et al.* have reported that deuterium incorporation at the gate loop remained unaffected upon ABA binding, suggesting that the gate loop remains flexible between open and closed conformations.^34^ Interestingly, we have observed large free energy barriers for ABA binding to both the monomeric and the dimeric PYL2 receptors, which are ~5 and ~4 kcal/mol respectively. The binding free energy barriers in PYL2 are comparable to those in both the PYL5 and the PYL10.^33^ Overall, our simulations show that ABA is required for PYL2 activation, and the dimeric PYL2 constrains the gate loop conformational plasticity and destabilizes the closed gate loop (active-like) conformation.

### Transition path theory characterizes the pathway of ABA binding and PYL2 activation

To obtain structural insights into PYL2 activation processes, we apply transition path theory (TPT) on the MSMs to determine the pathway associated with ABA binding and the conformational changes in both the monomeric and the dimeric receptors (Supplementary Methods).^53,54^ TPT characterizes the transition pathways in the MSMs that connect through ABA-unbound inactive MSM states and ABA-bound active MSM states, along with quantitative measure of the relative probabilities of the pathways.^53,54^ We report the top favorable pathway for ABA binding and PYL2 activation. The five transition states as captured in the top pathway are mapped onto the free energy landscape and labelled as states 1-5 (Fig. 2A, B). The snapshots of these states are shown in Fig. 2C and Fig. S3C.

Our results suggest that ABA binds to both monomeric and dimeric PYL2 receptors through a series of similar transition states, despite minor differences due to dimerization. The initial interaction between PYL2 and ABA is mediated by a direct contact between R120 and the carboxylate group of ABA, while the gate loop is open (state 1, Fig. 2C). Next, ABA enters into top of the binding pocket and interacts with both protomers, resulting in the stable intermediate state 2 (Fig. 2B). Specifically, the carboxylate group of ABA forms hydrogen bonding interactions with the side chains of R120 and S89 in one protomer, while the carbonyl group of ABA interacts with the backbone of Q90’ in the other subunit (state 2, Fig. 2C). Furthermore, the ring of ABA is sandwiched by F165 and L91’, which stabilizes ABA via *π-π* and hydrophobic interactions. We note that the interactions of ABA with Q90’ and L91’ can only occur in the dimeric PYL2 but not in the monomeric PYL2. Then, the carboxylate group of ABA moves towards the binding site to reach the intermediate state 3. At the state 3 (Fig. 2C), the carboxylate group and the hydroxyl group of ABA contact with the side chain of H119 and the backbone of I88, in addition to the hydrophobic interactions between ABA and several residues (F66, R120, F165, L91’). Both the gate loops remain open even after ABA reaches the binding site and forms a hydrogen bond with K64 (state 4, Fig. 2C). Finally, the gate loop in the ABA-bound protomer closes, while the gate loop in the ABA-free protomer remains open (state 5, Fig. 2C). In a previous study on PYL2 receptor^28^, the mutations of the residues in the binding pocket (K64A, F66A, F165A), the gate (S89A, L91A) and the latch (H119, R120) loops were suggested to result in significantly attenuated *in vitro* PYL2 activity on PP2C inhibition. The ABA binding pathway identified from our MD simulations and MSMs analysis highlights the critical roles of these mutated residues in ABA binding, in agreement with the mutation results.

Despite the similar activation pathways of both receptors, we note that there are key differences in relative stability of several transition states which depend on PYL2 oligomeric state (Fig. 2A, B). Notably, the intermediate state 3 is more stable in the dimeric PYL2 while the ABA-bound, closed gate loop state 5 is much less stable, compared to the monomeric PYL2. The free energy of state 3 in the dimeric PYL2 is ~2 kcal/mol, which is more stable (at least 1 kcal/mol in free energy difference) compared to that in the monomeric PYL2. The gain in stability is attributed to the dimer structure, where both protomers of PYL2 interact with ABA. In contrast, the state 3 in the monomeric PYL2 is less stable due to the weakened hydrophobic interaction between PYL2 and ABA (Fig. S3C). The state 5 in the dimeric PYL2 is less stable, which suggests that the dimerization makes PYL2 less likely to adopt the closed gate loop, active-like conformation.

The binding poses of ABA and the closed gate loop conformations predicted from our simulations deviate from the active PYL2 crystal structure less than 0.5 Å (PDB: 3KB3^28^) (Fig. S3D). Furthermore, we have captured the similar water-mediated interaction networks as in the crystal structure (Fig. S3E, F). Overall, our simulations elucidate the complete pathway of ABA binding and conformational change in PYL2, and suggest that the formation of dimeric complex stabilizes the binding intermediate state.

### Dimeric PYL2 stabilizes open gate loop conformation and lowers ABA binding affinity

To quantitatively characterize how ABA changes the populations of open and closed gate loop conformations, we classify the high-resolution MSM microstates into 4 different macrostates based on whether ABA is bound or unbound (4 Å as the cutoff PYL2-ABA distance) and whether the gate loop is open or closed (3 Å as the cutoff gate loop RMSD). The four-macrostate MSMs are then constructed to estimate the equilibrium probabilities of the macrostates and the transition probabilities between these macrostates (Fig. 3A, C). When ABA is not bound, the gate loop dominantly adopts an open conformation for both the monomeric and the dimeric PYL2. The probabilities of observing a closed gate loop conformation are 0.08% and 0.56% for the dimeric and the monomeric PYL2, respectively. The small population of closed gate loop states for the monomeric PYL2 may induce weak constitutive activity as reported in the previous experimental study. ^26^ Upon ABA binding, the binding of ABA enhances the equilibrium populations of closed gate loop states. The probabilities of observing a closed gate loop conformation are 7.4% and 20.4% for the dimeric and the monomeric PYL2, respectively. This suggests that the open gate loop conformation is more stable in the dimeric PYL2 relative to the monomeric PYL2. Overall, the binding of ABA is necessary for initializing the activation of PYL2 receptor, while the dimerization stabilizes the open gate loop conformation.

**Fig. 3.**
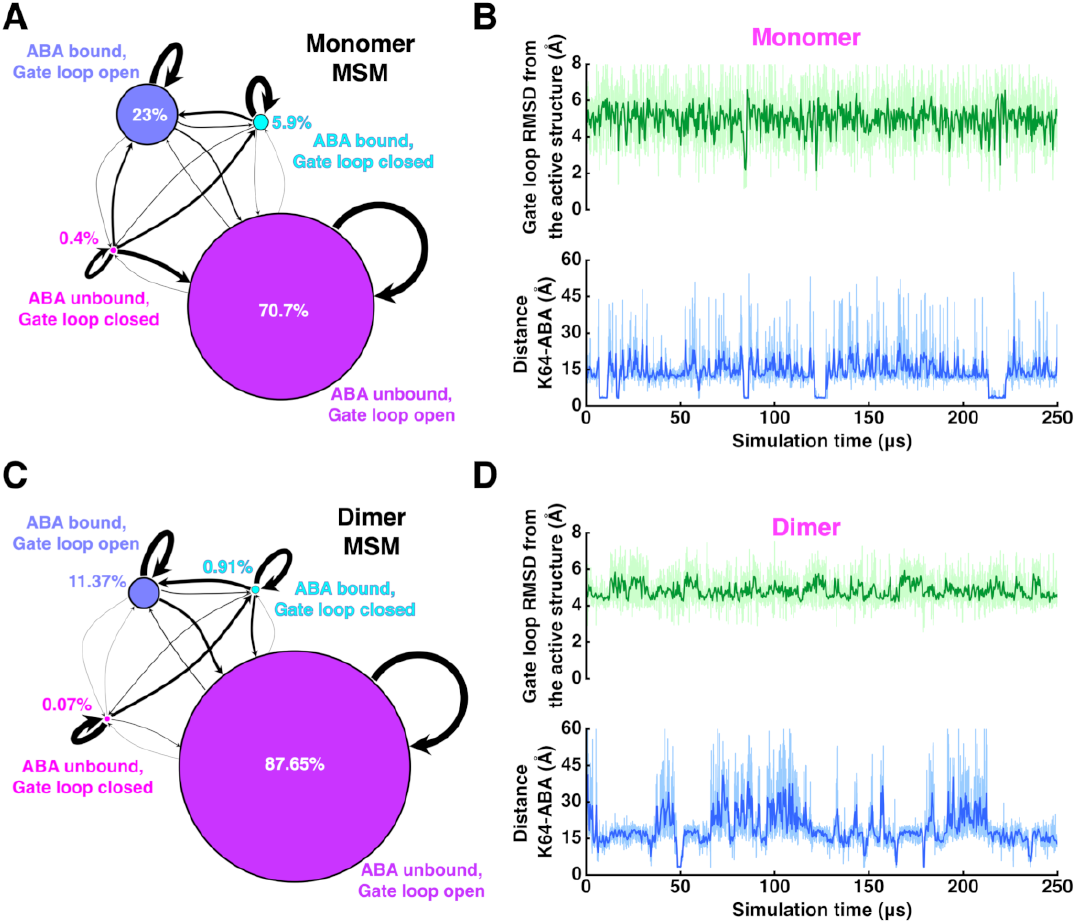
Kinetics of ABA binding to the monomeric and the dimeric PYL2 receptors. The conformational networks describe the ABA-mediated activation processes for (A) the monomeric and (C) the dimeric receptors. The four-macrostate (i.e. ABA unbound and gate loop open state, ABA unbound and gate loop closed state, ABA bound and gate loop open state as well as ABA bound and gate loop closed state) MSMs are constructed to represent the conformational space. The circles represent the macrostates in the MSMs, scaled by their MSM probabilities. The black edges represent the transitions between these macrostates, scaled by the square roots of their transitional probabilities. The macrostates are assigned from high-resolution MSMs according to the cutoff values for the PYL2-ABA distance and the gate loop RMSD from the activate structure. Long timescale dynamics of (B) the monomeric and (D) the dimeric PYL2 revealed by the conformational networks, showing the ABA binding and unbinding events (blue), the gate loop fluctuations (green) and the timescales at which they occur.

Next, we seek to characterize the long-timescale dynamics of monomeric and dimeric PYL2. From kinetic Monte Carlo simulations^55^ on the high-resolution MSMs, we generate 250 *μ*s long trajectories staring from an arbitrary ABA-unbound, open gate loop state (Supplementary Methods). For both the receptors, we have captured multiple ABA binding and unbinding events, and the associated conformational changes in the gate loop (Fig. 3B, D). It is apparent that the fluctuations in the gate loop conformation are significantly reduced in the dimeric PYL2 compared to the monomeric PYL2. When ABA is unbound, the gate loop RMSD primarily fluctuates between 3-8 Å for monomeric PYL2, and 4-7 Å for dimeric PYL2. When ABA is bound, for monomeric PYL2, the gate loop RMSD fluctuates between 1-8 Å suggesting that the gate loop closure is promoted by ABA binding but the loop remains flexible between open and closed conformations. However, for dimeric PYL2, we rarely observe that the gate loop RMSD reaches below 2 Å even after ABA is bound, suggesting that the stability of closed gate loop conformation decreases due to dimerization. We also note that the residence time of ABA in the bound state is substantially reduced by an order of magnitude for dimeric PYL2. In other words, ABA-bound pose is relatively less stable in dimeric PYL2 compared to monomeric PYL2. Thus, we conclude that the dimeric PYL2 stabilizes the open gate loop conformation and thereby lowers ABA binding affinity.

### Binding of ABA and subsequent closure of the gate loop destabilizes PYL2 complex

To quantitatively characterize how ABA changes thermostability of PYL2 complex, we seek to determine how standard PYL2 association free energy 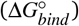 changes before and after ABA binding. We compare 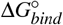 for the *apo-apo*, the *holo-apo*, and the *holo-holo* PYL2 complexes, where *apo* denotes an inactive protomer without ABA and *holo* denotes an active protomer with ABA bound. We employ potential of mean force (PMF)-based method for accurate estimation of 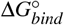.^44-46^ In each case, we separate two protomers along a vector *r* that connects the center of mass of two protomers in the presence of a series of geometrical restraints (Fig. 4A). To determine the separation PMF, we select a range of complex conformations with *r* evenly distributed in a certain range from the associated state to the fully dissociated state. Each conformation is used to seed a new replica simulation, with a harmonic potential acting on *r* to restrain the distance between two protomers. From these simulations, the separation PMF can be estimated via a well-established statistical free energy method, multistate Bennett acceptance ratio (MBAR). ^43^ The geometrical restraints are used to accelerate the convergence of separation PMF, which include the conformational restraints (*B*_1,*c*_, *B*_2,*c*_, *B*_1,*res*_, *B*_2,*res*_) and the restraints on the relative positions (*ϕ, θ*) and orientations (Θ, Φ and *ψ*) between two protomers (Fig. 4A). The contributions of separation PMF and all restraints to 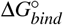 can be determined individually by integrating over their respective PMFs (Supplementary Methods).

**Fig. 4.**
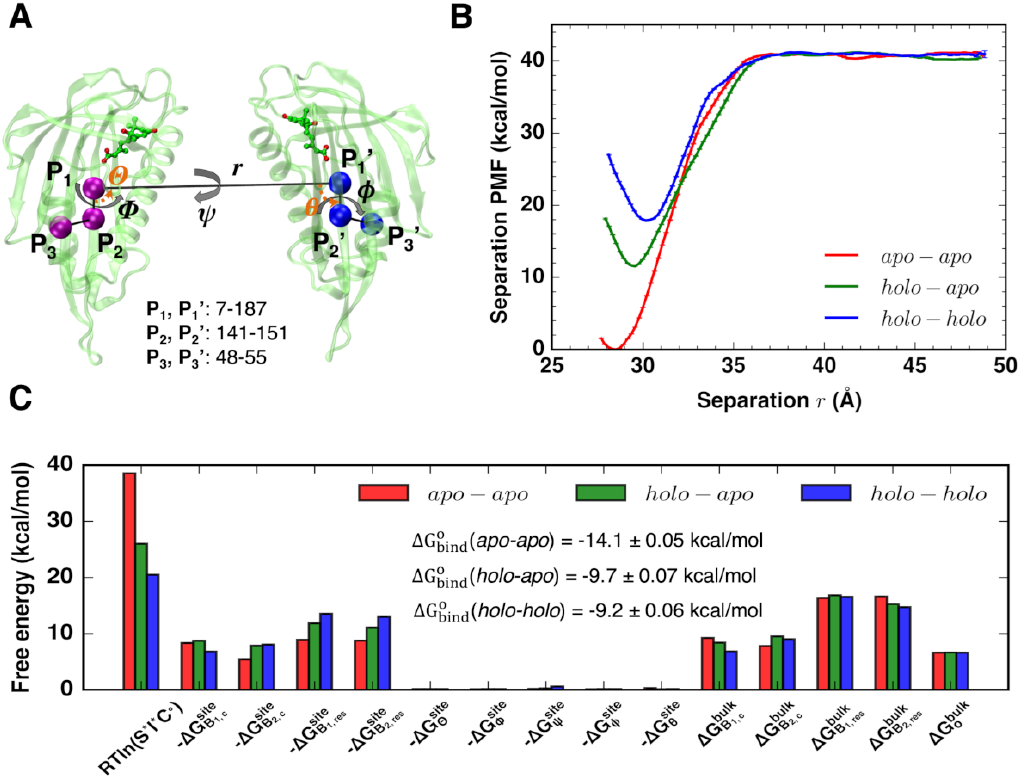
Determination of PYL2 homodimeric association free energies. (A) Snapshot of the *holo-holo* PYL2 and the collective variables used in PYL2 separation REUS simulations. The center of mass distance between the two protomers *r* (P_1_-P′_1_ distance), and the Euler angles *ϕ* (P_1_-P′_1_-P′_2_-P′_3_) and *θ* (P_1_-P′_1_-P′_2_) defines the relative position of the two protomers. The Euler angles, Θ (P′_1_-P_1_-P_2_), Φ (P′_1_-P_1_-P_2_-P_3_), and *ψ* (P′_2_-P′_1_-P_1_-P_2_), relate the relative orientation between the two protomers. In addition, PYL2 conformations are restrainted by RMSDs of the two protomers (*B*_1,*c*_, *B*_2,*c*_) and the interfacial residues (*B*_1,*res*_, *B*_2,*res*_) from the initial strutcures. (B) Potential of mean force (PMF) profiles for the separations of the *apo-apo* (red), the *holo-apo* (green), and the *holo-holo* (blue) PYL2. The error bars on the PMFs are shown. (C) Free energies associated to the components of 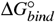 for the *apo-apo* (red), the *holo-apo* (green) and the *holo-holo* (blue) PYL2 complexes. The error bars for all components are less than 0.04 kcal/mol.

The starting structures of the *holo-apo*, and the *holo-holo* PYL2 complexes are obtained via short targeted MD simulations. We select 41 windows for each complex, with the distance *r* evenly distributed between 28-48 Å. We perform 8 ns replica exchange umbrella sampling (REUS) MD simulations for each replica, and use MBAR to determine the PMFs *W* (*r*). The separation PMFs for all three receptors are converged within 8 ns/window simulations (Fig. S4). The separation PMF profiles for the *apo-apo*, the *holo-apo*, and the *holo-holo* PYL2 complexes are given in Fig. 4B, along with the associated error bars. The wells in the three PMFs between 28-31 Å correspond to the associated states. *W* (*r*) converges to constant when *r* is greater than 38 Å, showing that the two protomers are fully separated. The well depths for the *apo-apo*, the *holo-apo*, and the *holo-holo* are 41 kcal/mol, 30 kcal/mol and 23 kcal/mol. The separation contributions to 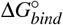 (denoted by *-RTln*(*I*S*C°*)) are determined by integrating over the separation PMFs and applying volume corrections, leading to −38.5 kcal/mol, −26.0 kcal/mol and −20.5 kcal/mol, respectively. It should be noted that these energy contributions include the effects of numerous restraints applied during the separation REUS MD simulations. However, these results qualitatively suggest that the stability of PYL2 complex decreases upon ABA binding. Also, we can observe that the minimum of *W*(*r*) slightly increases from 28.5 Å to 30 Å.

The contributions of adding the geometrical restraints in the associated state (denoted by site) and removing these restraints in the fully separated state (denoted by bulk) are calculated by determining PMFs for individual restraints (Supplementary Methods). In total, 8 PMFs for RMSD restraints acting on both the associated state and the separated state 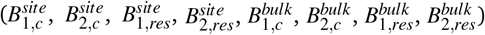 and 5 PMFs for angular restraints (*θ, φ, ψ, Φ, Θ*) acting on the associated state are determined via REUS MD simulations (Fig. S5). The convergence of all PMFs related to these restraints with respect to simulation time are carefully examined, and most PMFs converge after 8 ns/replica. For some PMFs related to RMSD restraints, we perform additional umbrella sampling simulations to expand the PMF regions and ensure the convergence. From these PMFs, we determine the contributions of all restraints to 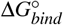 (Fig. 4C). Notably, the conformational restraints have relative large contributions to 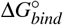, whereas the positional and orientational restraints have much smaller contributions.

Take together all individual contributions to 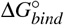, we obtain 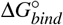 for the *apo-apo*, the *holo-apo*, and the *holo-holo* PYL2 as −14.1 kcal/mol, −9.7 kcal/mol and −9.2 kcal/mol, with the error bars less than 0.1 kcal/mol. The small error bars on 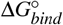 are due to the small error bars on PMFs. Our free energy calculation results show that the activation of one PYL2 protomer by ABA can introduce +4.4 kcal/mol difference in 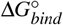, while activation of both protomers only further introduces +0.5 kcal/mol difference. The stability of PYL2 complexes thus decreases in response to ABA binding and receptor activation. However, 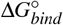 for both the *holo-apo* and the *holo-holo* PYL2 are fairly small, suggesting that PYL2 complex remains stable in the presence of ABA. These computational results are in agreement with previous experimental observations that PYL2 remains in dimer conformation both in the presence and in the absence of ABA.^27,29^ It is possible that PP2Cs might interact with the ABA-bound PYL2 in solution and contribute to the dissociation of PYL2. Overall, our free energy calculations quantitatively show that the binding of ABA destabilizes PYL2 complex by shifting the conformation equilibrium, while PYL2 remains stable after ABA binding.

### Presence of ABA substantially stabilizes PYL2-HAB1 complex and further promotes PYL2 dissociation

In the presence of both ABA and PP2C, ABA-bound PYL2 can interact with both PYL2 to form a stable homodimer and PP2C to form a heterodimer. We seek to determine 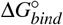 for PYL2-HAB1 complex to understand the competition between PYL2 homodimeric association and PYL2-PP2C heterodimeric association. We determine 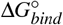 for both PYL2-HAB1 (in the absence of ABA) and PYL2-ABA-HAB1 (in the presence of ABA) complexes. Crystallographic studies have shown that PP2C form water-mediated interaction with ABA, and PP2C is therefore widely considered as a ABA co-receptor. ^32^ Biologically, it is also interesting to investigate how much the presence of ABA can stabilize PYL2-HAB1 complex.

Using a similar protocol as in PYL2 separation, the separation PMFs for the PYL2-HAB1 and the PYL2-ABA-HAB1 complexes are determined in the presence of conformational and angular restraints (Fig. 5A and Fig. S6A, B). The wells in the two PMFs between 39-41 Å correspond to the associated PYL2-HAB1 and PYL2-ABA-HAB1 complexes (Fig. 5B). The well depths for the PYL2-HAB1 and the PYL2-ABA-HAB1 are 28 kcal/mol and 33 kcal/mol, and their contributions to 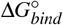 (−*RTln*(*I*S*C°*)) are −26.3 kcal/mol and −30.0 kcal/mol, respectively. These results suggest that the presence of ABA in the active PYL2 stabilizes the binding with HAB1. The contributions of adding the geometrical restraints in the site and removing them in the bulk are calculated by the PMFs for individual restraints (Fig. S6C, D and Fig. 5C). 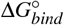 for PYL2-HAB1 and PYL2-ABA-HAB1 complexes are −10.5 kcal/mol and −18.3 kcal/mol, respectively. Our results show that the presence of ABA in PYL2 substantially stabilizes PYL2-HAB1 complex by −7.8 kcal/mol. The ABA’s stabilizing effects may be attributed to two aspects. First, the water-mediated interaction between ABA and HAB1 is expected to result in stronger association between PYL2 and HAB1. Second, the binding of ABA may perturb the conformational entropy of PYL2 receptor in the bulk and site, thereby contributing to the PYL2-HAB1 binding free energy. Overall, our results suggest that ABA is essential in stabilizing PYL2-HAB1 interactions.

**Fig. 5.**
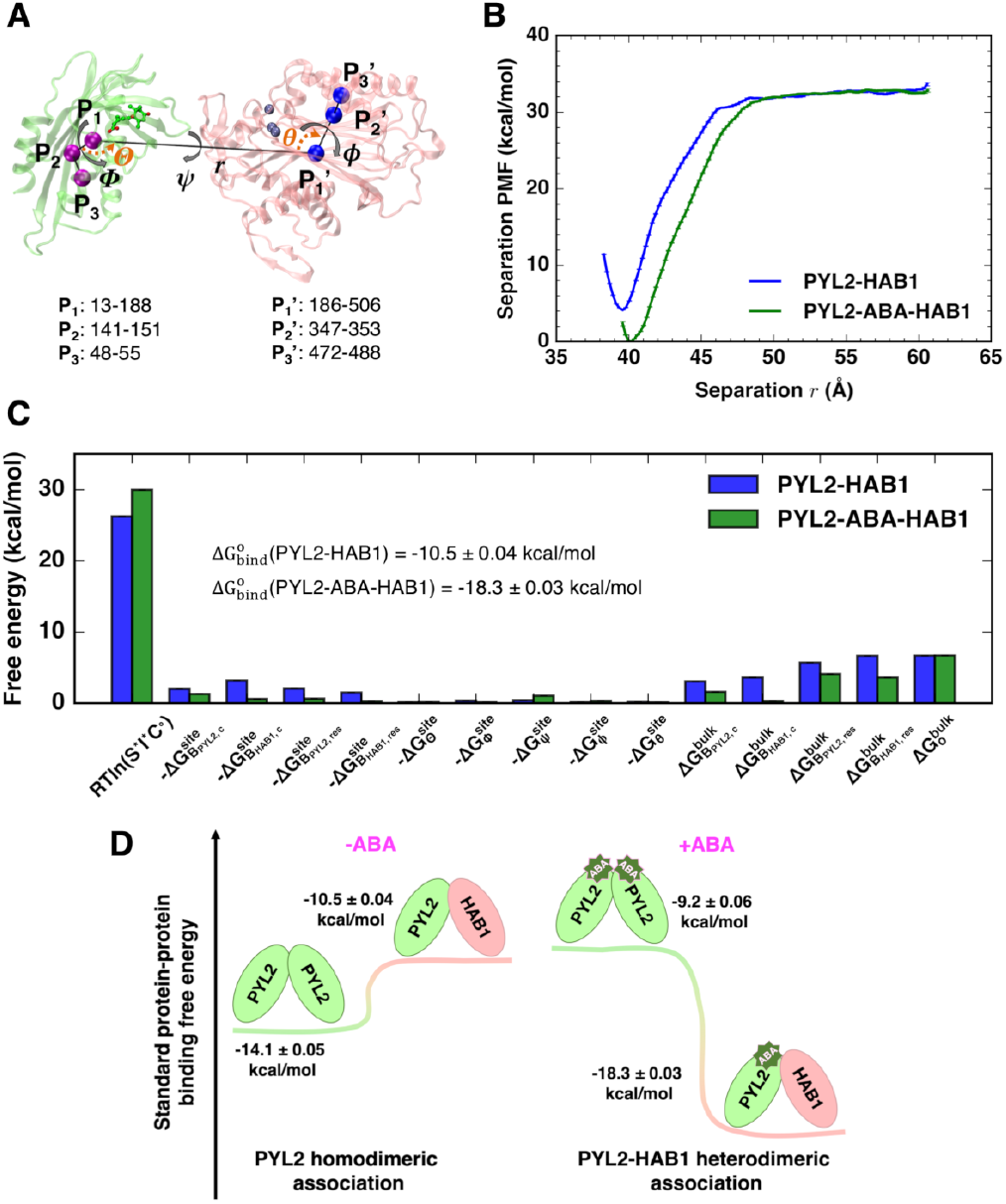
Determination of PYL2-HAB1 heterodimeric association free energies. (A) Snapshot of PYL2-ABA-HAB1 complex and the collective variables (*r*, *B*_*PYL*2,*c*_, *B*_*HAB*1,*c*_, *B*_*PYL*2,*res*_, *B*_*HAB*1,*res*_, Θ, Φ, *ψ, ϕ, θ*) used in separation REUS MD simulations. (B) PMFs for the separations of PYL2-HAB1 (red) and PYL2-ABA-HAB1 (blue) complexes, along with error bars on the PMFs. (C) Free energies associated to the components of 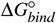 for PYL2-HAB1 and PYL2-ABA-HAB1 complexes, with error bars less than 0.03 kcal/mol. (D) Thermodynamic model for the competition between PYL2 homodimeric association and PYL2-HAB1 heterodimeric association in the absence and in the presence of ABA.

Take together 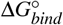 for PYL2 homodimeric association, we obtain a complete thermodynamic model to understand the competition between PYL2 homodimeric association and PYL2-HAB1 heterodimeric association (Fig. 5D). In the absence of ABA, 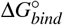 for *apo-apo* PYL2 and PYL2-HAB1 complex are −14.1 kcal/mol and −10.5 kcal/mol, respectively. PYL2 homodimeric association is favored compared to heterodimeric association between PYL2 and PP2C. In the presence of ABA, PYL2 complex is destabilized and PYL2-HAB1 complex is stabilized by the binding of ABA. 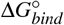 for *holo-holo* PYL2 and PYL2-ABA-HAB1 complex are −9.2 kcal/mol and −18.3 kcal/mol. As a result, heterodimeric association between PYL2 and HAB1 is largely favored than PYL2 homodimeric association. Overall, our results suggest that the presence of ABA stabilizes PYL2-HAB1 complex and further promotes PYL2 dissociation.

## Conclusions

In this study, we have investigated the atomic scale dynamics of ABA perception by PYL2 in both its monomeric and dimeric forms, allowing for a detailed understanding of how dimerization affects receptor activation and ABA binding affinity. In addition, we have characterized how ABA binding affects PYL2 dissociation and PYL2-HAB1 association processes. On ABA perception, our MD simulations have uncovered the key intermediate states involved in the binding of ABA and subsequent activation of both monomeric and dimeric PYL2. The key residues that mediate PYL2-ABA interactions in our predicted intermediate states have been shown to be critical for ABA receptor activity from previous mutation studies. ^28^ Free energy landscapes suggest that PYL2 complex substantially constrains the conformational plasticity of the gate loop, thereby stabilizing the open conformational state and destabilizing the closed state. Furthermore, the PYL2 complex stabilizes the intermediate state due to the enhanced hydrophobic interactions. As a result, the ABA off-binding rate increases due to dimerization, leading to lower ABA affinity in the dimeric PYL2. On receptor activation, our results suggest that ABA is required to trigger the gate loop closure for the dimeric PYL2, and the gate loop remains in open conformation even after ABA binding, in agreement with HDX experimental data.^34^ In this sense, ABA binding is necessary but far from sufficient condition for the activation of the dimeric PYL2. Overall, our simulations have explained the molecular basis of intrinsic lower ABA sensitivity for dimeric ABA receptors.

On PYL2 dissociation, our free energy calculations suggest that the activation of both protomers introduce ∼4.9 kcal/mol differences in the stability of PYL2 complex. However, both the *holo-apo* and the *holo-holo* complexes are highly stable, with free energies of −9.7 kcal/mol and −9.2 kcal/mol, respectively. The binding of ABA can only slightly shift the equilibrium of PYL2 complex formation. The PYL2 dissociation is further promoted by the presence of PP2C in solution. It should be noted that ABA-bound PYL2 interacts with PP2Cs in solution with a dissociation constant *K_d_* between 150-250 nM.^28,29^ Our free energy calculation results show that the presence of ABA substantially decrease PYL2-HAB1 association free energy from −10.5 kcal/mol to −18.3 kcal/mol. Therefore, the presence of ABA makes the homodimeric PYL2 association less competitive than the heterodimeric PYL2-PP2Cs association as compared to that in the absence of ABA. Overall, our thermodynamic model explains the competitive associations between PYL2 homodimerization and PYL2-HAB1 heterodimerization.

In *Arabidopsis thaliana*, PYL2, PYL5 and PYL10 receptors belong to three individual classes of PYR/PYL/RCAR receptors based on their sequence similarities. ^16^ Combining the results from our recent work on PYL5 and PYL10,^33^ we can draw several conclusions regarding the similarities and the differences in activation of the three classes of ABA receptors. On the similarities, (1) ABA binding is necessary but not sufficient for full activation of PYR/PYL/RCAR receptors. We have shown that the binding of ABA can shift the conformational equilibrium of the gate loop, thereby increasing the equilibrium probability of closed gate loop conformations. However, the gate loop remains flexible between open and closed conformations. ^34^ Strikingly, the dimerization of PYR/PYL/RCAR receptors substantially favor the open gate loop conformations, due to the fact that the gate loop is involved in forming the dimer interface. PP2C binding is required to stabilize the closed gate loop conformations for full receptor activation. In the case of dimeric receptors, PP2C might be required for their full dissociation. (2) A large binding free energy barrier (∼4-5 kcal/mol) is observed for all the receptors, which is associated with the exclusion of water molecules from the binding pocket.^33^ The displacement and reorganization of water networks in the binding site may contribute to overall free energy of ABA binding.

On the differences among three clades of ABA receptors in *Arabidopsis thaliana*, (1) the ABA binding pathways and the relative stability of transition states in the pathways vary depending on the sequence and the oligomeric state. While the key polar residues that mediate ABA binding are conserved, the binding pathways are not conserved across ABA receptors. For example, we previous reported that the side chain switch of F58 in PYL10 is required for ABA binding, ^33^ whereas this is not the same in both PYL2 and PYL5 receptors. In this study, we have shown that PYL2 complex stabilizes the binding intermediate state while destabilizes the active state. It is possible that the variations in solvation structural and thermodynamic properties across different ABA receptors due to amino acid substitutions contribute to the binding differences. (2) ABA-independent and ABA-dependent activation depends on the sequence and the oligomeric state. For the monomeric receptors, we have shown that the gate loop has a higher degree of conformational plasticity, which results in the experimentally observed ABA-independent activity of these receptors. Notably, the PYL10 has shown a distinct constitutive activity.^26,30^ For the dimeric receptors, ABA binding is required to trigger receptor activity because the dimer conformation cannot be broken by PP2Cs in the absence of ABA.

Finally, recent work from our group has tried to provide a comprehensive structural and dynamic view of plant hormone signaling for ABA^33^, Brassinosteroid^40,56,57^ and Strigolactone^58^ hormone receptors in plants. These physical insights into the structural, dynamic and energetic basis of the activation of hormone receptors in *Arabidopsis thaliana*, which are challenging to obtain with experimental approaches. Beyond *Arabidopsis thaliana*, there are a growing number of studies on hormone receptors in higher plants, such as rice^59^, soybean^60^ and maize^61^. Our key findings on the activation mechanism of ABA and other hormone receptors could be generalized across other plant species given that several core components of hormone signaling pathway and the sequences of hormone receptors are highly conserved. However, the structural information on hormone receptors in higher plants is still scarce^62,63^. Therefore, there is a need to develop computational approaches that could reliably predict protein structures from sequence, allow rapid conformational sampling and experimental characterization of the hormone receptors in plants^50,51,64,65^.

## Conflicts of interest

There are no conflicts to declare.

## Acknowledgements

The authors acknowledge the support from the Blue Waters sustained-petascale computing project, which is funded by the National Science Foundation (awards OCI-0725070 and ACI-1238993) and the state of Illinois. D.S. acknowledges the support from Foundation for Food and Agriculture Research via the New Innovator Award in Food & Agriculture Research. C.Z. acknowledges the support by 3M Corporate Fellowship and Ullyot Graduate Fellowship from the University of Illinois. The authors thank Alexander S. Moffett from the University of Illinois for the helpful discussions on standard protein-protein binding free energy calculations.

## Author contributions

C.Z. and D.S. designed the project. C.Z. performed simulations, analyzed data, and wrote manuscript with input from D.S.

## Notes

### Competing Interest Statement

The authors have declared no competing interest.

### Summary of Updates

This version is a revised manuscript based on the reviewer comments from Chemical Science.

