## Supplementary File for "Molecular Basis of the Activation and Dissociation of Dimeric PYL2 Receptor in Abscisic Acid Signaling"

### Electronic Supplementary Information (ESI)

### Methods

#### General MD Simulation Details

The computational approaches and software used for different aspects of this study were summarized in Table S1. All MD simulations were set up using AmberTools15 and performed using Amber[1] and NAMD[2] with the AMBER ff14SB force field for protein and the general AMBER force field[3] for ABA. NAMD was used in replica exchange umbrella sampling (REUS) MD simulations for standard protein-protein binding free energy calculations, due to the ease of definition of collective variables used for running REUS MD simulations. Amber was used for other simulations for computational efficiency. The partial charges of ABA were determined according to AM1-BCC method using Antechamber in the AmberTools15. The protein and ABA were solvated with explicit TIP3P waters. The simulation box was neutralized by adding counterions $Na^+$ and $Cl^-$. Extra counterions were added to create an ionic concentration of 0.15 M. The SHAKE algorithm[4] was applied to constrain the length of covalent bonds involving hydrogen atoms. The particle-mesh Ewald method[5] was used to treat electrostatic interactions, with a 10 Å cutoff for van der Waals interactions. Production simulations were run on Blue Waters supercomputer with an integration time step of 2 fs.

#### MD Simulations and Adaptive Sampling of ABA Binding to the Monomeric and the Dimeric PYL2 Receptors

The crystal structure of inactive PYL2 receptor (PDB: 3KDH[6]) was used as the starting structure for MD simulations of ABA binding processes. The full dimer structure (chains A and B) was used for the dimer simulations and the chain A was used for the monomer simulations. Single ABA molecule was randomly placed away from the binding pocket using Packmol[7]. The receptor-ABA complexes were then solvated with a large water box that exceeds complex surface by at least 10 Å. After 10000 steps minimization, the system was heated to 300 K and equilibrated in isothermal-isobaric (NPT, 300 K, 1 atm) ensemble for 1 ns. Production runs were then launched without applying any artificial potential.

Parallel MD simulations were initialized and continued with multiple rounds of adaptive sampling[8,9,10,11] to achieve ABA binding and subsequent gate loop closure (Tables S2 and S3). The adaptive sampling scheme has been employed in several published studies on protein-ligand binding simulations.[12,13,14] For the first round of simulations, the ligands were randomly positioned at multiple positions outside of the binding site of PYL2 receptor. For the next round of simulations, the starting configurations were chosen from the simulation snapshots which have the ligand at the minimum distance from the binding pocket. The simulations were stopped when we have observed ligand binding and obtained sufficient simulation data to build a Markov State Model of the ligand binding process. For each round of adaptive simulations, the initial velocities of all atoms were randomly generated from Maxwell-Boltzmann distribution to start new simulations. Amber14 and Amber18 were used for the dimeric and the monomeric PYL2 systems, respectively. The aggregate simulation times for the dimeric and the monomeric PYL2 were ~116 $\mu$s and ~107 $\mu$s, respectively.

#### Featurization of Binding MD Trajectories

The ABA binding MD simulation datasets were featurized using MDTraj 1.7.0[15]. This featurization step is to calculate a series of structural distance metrics that are used for clustering all the conformations collected from MD simulations. A similar set of features as used in our previous work[16], including the 32 atomic distances between PYL2 and ABA, were employed to analyze the monomeric PYL2 simulations (Table S4). These distances characterize the conformational changes in both the gate loop and the latch loop, the movement of ABA, and the molecular interactions between PYL2 and ABA. For the dimeric PYL2, the sets of distances between ABA and both protomers were calculated, leading to 64 distances in total for clustering analysis.

#### Construction and Optimization of Markov State Models

Time-lagged independent component analysis (tICA) was performed on these featurization metrics in order to capture several slowest-relaxing degrees of freedom (referred to as tICs) from linear combinations of the distance features[17]. All conformations collected from MD simulations were then clustered into $N$ states based on $M$ slowest tICs using the

$k$-means clustering method. Markov state models (MSMs) were then constructed based on the clustering results. The matrix of transition probabilities was determined with a lag time $\tau$ using maximum likelihood approximation. Similar to our previous work[16], the optimal lag times ($\tau$) were chosen based on the convergence of the implied timescales of MSMs and the number of clusters ($N$) as well as the number of tICs ($M$) were optimized via cross validation ranked by the variational GMRQ objective function[18] (Fig. S7). The MSM parameters ($\tau$, $N$, $M$) for the monomeric PYL2 are 40 ns, 300 clusters and 4 tICs and for the dimeric PYL2 are 40 ns, 200 clusters and 9 tICs. All MSMs were constructed using the MSMBuilder 3.4[19] and the MSM hyperparameters were optimized using the Osprey software[20].

#### Kinetic Monte Carlo Simulations

Kinetic MC simulations[21] were performed on the built MSMs to predict the long-timescale dynamics of the monomeric and the dimeric PYL2 receptors associated with ABA binding. The kinetic MC algorithm is similar to a random walk process on the receptor conformational networks, where the probabilities of jumping between different states in the network are given by the MSM transition probability matrix. All the kinetic MC simulations were performed using the MSMBuilder 3.4[19]. The state with the largest K64-ABA distance was chosen as the arbitrary starting state.

#### Transition Path Theory

Transition path theory (TPT) is a numerical method for analyzing rare events in complex dynamical systems[22,23]. Applying TPT in kinetic network models allows for characterization of reactive probabilities and fluxes of the pathways captured by the MSMs. In this case, the pathways between the ABA-unbound inactive states and the ABA-bound active states with the highest fluxes can be identified from TPT. All the analysis was implemented using MSMBuilder 3.4[19].

#### Calculation of Standard Binding Free Energy for Protein-Protein Association

The potential of mean force (PMF)-based method for accurate estimation of standard protein-protein binding free energy were established from the previous studies[24,25,26]. The absolute protein-protein association free energy ($\Delta G^o_{bind}$) was determined by calculating PMF for separating two proteins in the presence of a series of conformational, positional and orientational restraints. These restraints substantially reduce the configuration entropy of complex and accelerate the convergence of separation PMF. Integration of the separation PMF contributes to the major component of $\Delta G^o_{bind}$. Next, the contributions of adding these restraints on the associated state and removing them from the fully dissociated state to $\Delta G^o_{bind}$ were also computed and then subtracted from the free energy component resulting from separation PMF. In total, there were 9 harmonic restraints applied during separation, including four relating to conformational changes in two proteins and five relating to relative orientations and positions of two proteins (Fig. 6A and Fig. 7A). The conformational restraints acting on heavy-atom RMSD of the backbone in each protein (denoted by subscripts $B_{1,c}$, $B_{2,c}$) and heavy-atom RMSD of the side chains in the interfacial residues (denoted by subscripts $B_{1,res}$, $B_{2,res}$) restrict protein structural fluctuations and deviations from the starting associate state. Three angular restraints acting on the relative orientation of two proteins (denoted by subscript $o$, including $\Theta$, $\Phi$, $\Psi$) and two on the relative position (denoted by $a$, including $\phi$, $\theta$) limit the configuration space when two proteins are separated. For each restrained collective variable (CV) $\xi_i$, a corresponding harmonic potential ($u_{\xi_i}$) is added to the system Hamiltonian, which is defined as follow:

$$u_{\xi_i} = \frac{1}{2} k_{force,i} (\xi_i - \xi_{i,ref})^2 \tag{1}$$

where $k_{force,i}$ is the force constant and $\xi_{i,ref}$ is the reference value for each CV.

Given the equilibrium association constant, $\Delta G^o_{bind}$ is given by

$$\Delta G^o_{bind} = -\beta^{-1} ln(K_{eq} C^o) \tag{2}$$

where $\beta$ is the reciprocal of the product of gas constant and temperature, and $C^o$ is standard concentration of 1 M, which is 1/1661 Å$^3$. $K_{eq}$ can be expressed as in equation 3.

$$
\begin{aligned}
K_{eq} = S^* I^* e^{-\beta[(G^{bulk}_{B_{1,c}} - G^{site}_{B_{1,c}}) + (G^{bulk}_{B_{2,c}} - G^{site}_{B_{2,c}})]} \\
* e^{-\beta[(G^{bulk}_{B_{1,res}} - G^{site}_{B_{1,res}}) + (G^{bulk}_{B_{2,res}} - G^{site}_{B_{2,res}})]} \\
* e^{-\beta[(G^{bulk}_{o} - G^{site}_{o}) - G^{site}_{a}]}
\end{aligned}
\tag{3}
$$

The term $S^*$ addresses the removal of positional restraints ($\theta$, $\phi$) on one protein, which is separated from the other protein along $r$ instead of free diffusion. $S^*$ is given by

$$
S^* = r^{*2} \int_0^{\pi} d\theta sin\theta \int_0^{2\pi} d\phi e^{-\beta u_a(\theta,\phi)} \tag{4}
$$

where $r^*$ is a point far from the binding site and $u_a = u_\theta + u_\phi$. The term $I^*$ is given by

$$
I^* = \int_{site} dr e^{-\beta[W(r) - W(r^*)]} \tag{5}
$$

which includes the contribution from the separation PMF $W(r)$.

The exponential terms in equation 3 are related to adding the restraints on the associated state (denoted by site) and removing the restraints from the dissociated state (denoted by bulk). Due to the configuration isotropy in the bulk, the term $G^{bulk}_o$ relating to removing the bulk orientational restraints can be calculated through direct numeric integration as follow:

$$
e^{-\beta G^{bulk}_o} = \frac{1}{8\pi^2} \int_0^{\pi} d\Theta \int_0^{2\pi} d\Phi \int_0^{2\pi} d\Psi sin\Theta e^{-\beta u_o(\Theta,\Phi,\Psi)} \tag{6}
$$

In order to determine the remaining terms, 8 terms involving the conformational restraints in both the bulk and the site ($G^{site}_{B_{1,c}}$, $G^{site}_{B_{2,c}}$, $G^{site}_{B_{1,res}}$, $G^{site}_{B_{2,res}}$, $G^{bulk}_{B_{1,c}}$, $G^{bulk}_{B_{2,c}}$, $G^{bulk}_{B_{1,res}}$, $G^{bulk}_{B_{2,res}}$) as well as 5 terms ($G^{site}_a$ and $G^{site}_o$) involving the orientational and positional restraints in the site, additional PMFs for each restraint need to be determined from MD simulations. The individual free energies are given as follow:

$$
e^{\beta G^{site}_{B_{1,c}}} = \frac{\int_{site} d\mathbf{1} \int \mathbf{dXe}^{-\beta U}}{\int_{site} d\mathbf{1} \int \mathbf{dXe}^{-\beta(U + u_{B_{1,c}})}} = < e^{\beta u_{B_{1,c}}} >_{(site,U)} \tag{7}
$$

$$
e^{\beta G^{site}_{B_{2,c}}} = \frac{\int_{site} d\mathbf{1} \int \mathbf{dXe}^{-\beta(U + u_{B_{1,c}})}}{\int_{site} d\mathbf{1} \int \mathbf{dXe}^{-\beta(U + u_{B_{1,c}} + u_{B_{2,c}})}} = < e^{\beta u_{B_{2,c}}} >_{(site,U,u_{B_{1,c}})} \tag{8}
$$

$$
e^{\beta G^{site}_{B_{1,res}}} = \frac{\int_{site} d\mathbf{1} \int \mathbf{dXe}^{-\beta(U + u_{B_{1,c}} + u_{B_{2,c}})}}{\int_{site} d\mathbf{1} \int \mathbf{dXe}^{-\beta(U + u_{B_{1,c}} + u_{B_{2,c}} + u_{B_{1,res}})}} = < e^{\beta u_{B_{1,res}}} >_{(site,U,u_{B_{1,c}},u_{B_{2,c}})} \tag{9}
$$

$$
e^{\beta G^{site}_{B_{2,res}}} = \frac{\int_{site} d\mathbf{1} \int \mathbf{dXe}^{-\beta(U + u_{B_{1,c}} + u_{B_{2,c}} + u_{B_{1,res}})}}{\int_{site} d\mathbf{1} \int \mathbf{dXe}^{-\beta(U + u_{B_{1,c}} + u_{B_{2,c}} + u_{B_{1,res}} + u_{B_{2,res}})}} = < e^{\beta u_{B_{2,res}}} >_{(site,U,u_{B_{1,c}},u_{B_{2,c}},u_{B_{1,res}})} \tag{10}
$$

$$
\begin{aligned}
e^{\beta G^{site}_{o}} = \frac{\int_{site} d\mathbf{1} \int \mathbf{dXe}^{-\beta(U + u_{B_{1,c}} + u_{B_{2,c}} + u_{B_{1,res}} + u_{B_{2,res}})}}{\int_{site} d\mathbf{1} \int \mathbf{dXe}^{-\beta(U + u_{B_{1,c}} + u_{B_{2,c}} + u_{B_{1,res}} + u_{B_{2,res}} + u_o)}} \\
= < e^{\beta u_o} >_{(site,U,u_{B_{1,c}},u_{B_{2,c}},u_{B_{1,res}},u_{B_{2,res}})}
\end{aligned}
\tag{11}
$$

$$e^{\beta G_a^{site}} = \frac{\int_{site} d\mathbf{1} \int \mathbf{dXe}^{-\beta(U+u_{B_{1,c}}+u_{B_{2,c}}+u_{B_{1,res}}+u_{B_{2,res}}+u_o)}}{\int_{site} d\mathbf{1} \int \mathbf{dXe}^{-\beta(U+u_{B_{1,c}}+u_{B_{2,c}}+u_{B_{1,res}}+u_{B_{2,res}}+u_o+u_a)}} =< e^{\beta u_a} >_{(site,U,u_{B_{1,c}},u_{B_{2,c}},u_{B_{1,res}},u_{B_{2,res}},u_o)} \tag{12}$$

$$e^{-\beta G_{B_{1,c}}^{bulk}} = \frac{\int_{bulk} d\mathbf{1}\delta(\mathbf{r_1}-\mathbf{r_1^*}) \int \mathbf{dXe}^{-\beta(U+u_{B_{1,c}})}}{\int_{bulk} d\mathbf{1}\delta(\mathbf{r_1}-\mathbf{r_1^*}) \int \mathbf{dXe}^{-\beta U}} =< e^{-\beta u_{B_{1,c}}} >_{(bulk,U)} \tag{13}$$

$$e^{-\beta G_{B_{2,c}}^{bulk}} = \frac{\int_{bulk} d\mathbf{1}\delta(\mathbf{r_1}-\mathbf{r_1^*}) \int \mathbf{dXe}^{-\beta(U+u_{B_{1,c}}+u_{B_{2,c}})}}{\int_{bulk} d\mathbf{1}\delta(\mathbf{r_1}-\mathbf{r_1^*}) \int \mathbf{dXe}^{-\beta(U+u_{B_{1,c}})}} =< e^{-\beta u_{B_{2,c}}} >_{(bulk,U,u_{B_{1,c}})} \tag{14}$$

$$e^{-\beta G_{B_{1,res}}^{bulk}} = \frac{\int_{bulk} d\mathbf{1}\delta(\mathbf{r_1}-\mathbf{r_1^*}) \int \mathbf{dXe}^{-\beta(U+u_{B_{1,c}}+u_{B_{2,c}}+u_{B_{1,res}})}}{\int_{bulk} d\mathbf{1}\delta(\mathbf{r_1}-\mathbf{r_1^*}) \int \mathbf{dXe}^{-\beta(U+u_{B_{1,c}}+u_{B_{2,c}})}} =< e^{-\beta u_{B_{1,res}}} >_{(bulk,U,u_{B_{1,c}},u_{B_{2,c}})} \tag{15}$$

$$e^{-\beta G_{B_{2,res}}^{bulk}} = \frac{\int_{bulk} d\mathbf{1}\delta(\mathbf{r_1}-\mathbf{r_1^*}) \int \mathbf{dXe}^{-\beta(U+u_{B_{1,c}}+u_{B_{2,c}}+u_{B_{1,res}}+u_{B_{2,res}})}}{\int_{bulk} d\mathbf{1}\delta(\mathbf{r_1}-\mathbf{r_1^*}) \int \mathbf{dXe}^{-\beta(U+u_{B_{1,c}}+u_{B_{2,c}}+u_{B_{1,res}})}} =< e^{-\beta u_{B_{1,res}}} >_{(bulk,U,u_{B_{1,c}},u_{B_{2,c}},u_{B_{1,res}})} \tag{16}$$

where $U$ is the net potential energy without any restraint. The expectation values in the equations 7-16 can be calculated through use of the PMF $W(\xi)$ of the relevant CV $\xi$ in the given ensemble. For example, the contribution due to the bulk conformational restraint ($B_{1,c}$) is given by the equation 17.

$$e^{-\beta G_{B_{1,c}}^{bulk}} =< e^{-\beta u_{B_{1,c}}} >_{(bulk,U)} = \frac{\int_{bulk} d\xi e^{-\beta W(\xi)} e^{-\beta u_{B_{1,c}}(\xi)}}{\int_{bulk} d\xi e^{-\beta W(\xi)}} \tag{17}$$

#### Targeted MD Simulations of the Dimeric PYL2 Receptor

Targeted MD simulations starting from the crystal structure of the *apo-apo* PYL2 (PDB: 3KDH[6]) were used to generate the *holo-apo* and the *holo-holo* PYL2 structures. Initially, one or two ABA were placed away from the *apo-apo* PYL2 crystal structure, respectively. The PYL2-ABA complexes were then solvated with explicit waters. After equilibration for 1 ns, a series of 0.5 ns targeted MD simulations were performed sequentially using Amber 18 to capture ABA binding and subsequent closures of the gate loop in one or both of the ABA-bound protomers. The PYL2 active crystal structure (PDB: 3KDI[6]) was used as the target structure. Intrinsically, targeted MD adds an additional harmonic potential to the system Hamiltonian, which is based on the root mean square deviations (RMSD) of ABA and the gate loop from the target structure. Due to the biasing potentials, ABA and the gate loop move toward the *holo-apo* and the *holo-holo* PYL2 structures within short amounts of MD simulations. The applied force constant was 30 kcal·mol$^{-1}$·Å$^{-2}$.

#### Replica Exchange Umbrella Sampling MD Simulations of PYL2 Separation and Determination of Separation PMF

Replica exchange umbrella sampling (REUS) MD simulations[27] were used to calculate the separation PMFs of the *apo-apo*, the *holo-apo*, and the *holo-holo* PYL2 in the presence of conformational, positional and orientational restraints. REUS simulations were performed using NAMD 2.13 software[2]. The *apo-apo* crystal structure, and the *holo-apo*, the *holo-holo* structures from targeted MD simulations were used as the staring structures. The complexes were solvated with a sufficiently large water box which encompasses both protomers when fully separated. After 20000 steps energy minimization, each system was equilibrated in NPT ensemble for 1 ns. An additional 200 ps simulation for each system was performed to measure the average values of all restrained CVs in the associated state. Next, steered MD simulations were performed to slowly increase the center of mass distance between the two protomers ($r$) to 48 Å over 8 ns, in order to generate starting structures for REUS MD simulations. From the trajectory, 41 structures with $r$ evenly spaced between 28 and 48 Å were chosen as the starting structures for REUS MD simulations (41 windows, 0.5 Å/window). The starting structure in each umbrella window was equilibrated for 1 ns and then each replica was run for 8 ns. The force constants and the reference values for the restrained CVs were given in Table S5.

The separation PMF was estimated using the multistate Bennett acceptance ratio (MBAR) method with pymbar python package[28]. The trajectories from REUS MD simulations were sorted and subsampled to ensure uncorrelated samples for estimating PMF. The integration of separation PMF $W(r)$ and the error bar on the integration $\sigma_{I^*}$ from error propagations are given by

$$I^* = \sum_{i=1}^{N_{site}} \Delta r_i e^{-\beta[W(r_i)-W(r^*)]} \tag{18}$$

$$\sigma_{I^*}^2 = \sum_{i=1}^{N_{site}} (\Delta r_i \beta e^{-\beta[W(r_i)-W(r^*)]})^2 \sigma_{W(r_i)}^2 \tag{19}$$

where $\sigma_{W(r_i)}$ is the error bar on the PMF reported by MBAR.

#### Replica Exchange Umbrella Sampling MD Simulations of PYL2-HAB1 Separation and Determination of Separation PMF

REUS MD simulations were performed to calculate the separation PMFs of the PYL2-HAB1 and the PYL2-ABA-HAB1 complexes using a similar protocol in PYL2 separation. The crystal structure of PYL2-ABA-HAB1 complex (PDB: 3KB3[29]) was used as the starting structure. The ABA molecule was removed to get the PYL2-HAB1 complex. Both the PYL2-HAB1 and the PYL2-ABA-HAB1 complexes were then solvated with a sufficiently large water box. The MC/MD simulation was performed for 2325 cycles (10,000 MC steps and 1,000 MD steps) to equilibrate water occupancy for the buried cavities in PYL2-HAB1 complex. Then, the equilibrated systems were subjected to steered MD simulations to generate initial structures for REUS MD simulations. In both cases, 41 structures with $r$ evenly spaced between 40 and 60 Å were chosen as the starting structures for REUS MD simulations (41 windows, 0.5 Å/window). The starting structure in each umbrella window was equilibrated for 1 ns and then each replica was run for 8 ns. The force constants and the reference values for the restrained CVs were given in Table S6. For the PYL2-HAB1 complex, additional targeted MD simulations were run to generate extra 4 windows spanning between 38 and 40 Å (0.5 Å/window) and umbrella sampling simulations were performed for 12 ns/window. The separation PMF was estimated using MBAR, and the integration of separation PMF and error propagations were determined using the equations 18 and 19.

#### Determination of PMFs for Individual Restrained Collective Variables

REUS MD simulations along the restrained CVs were performed to calculate the free energy contributions of adding those restraints to the associated state and removing them from the dissociated state. To generate a wide range of starting structures for each restrained CV, we performed 6 ns temperature-accelerated MD simulations, coupling with the respective CV to a dummy particle experiencing a temperature of 2500 K. The starting structures for REUS windows were chosen from the accelerated MD trajectory, with a window size of 0.05 Å for conformational restraints and 1° for angular restraints. Each replica was equilibrated for 1 ns and then run for 8 ns. The force constants used are 1000 kcal·mol$^{-1}$·Å$^{-2}$ for RMSD restraints and 2.5 kcal·mol$^{-1}$·$^{\circ -2}$ for angular restraints. The PMFs were determined using

MBAR. For some of the RMSD restraints, we performed additional targeted MD simulations and umbrella sampling simulations to expand the ranges of PMFs and ensure the convergence of PMFs.

Given the PMFs, the expectation values in the equation 7-16 and the associated error bars can be calculated via numerical computations. For example, $< e^{-\beta u_{B_{1,c}}} >_{(bulk,U)}$ (denoted by $A$) is given by

$$e^{-\beta G^{bulk}_{B_{1,c}}} =< e^{-\beta u_{B_{1,c}}} >_{(bulk,U)}= A = \frac{\sum_{i=1}^{N_{bulk}} \Delta\xi_i e^{-\beta W(\xi_i)} e^{-\beta u_{B_{1,c}}(\xi_i)}}{\sum_{i=1}^{N_{bulk}} \Delta\xi_i e^{-\beta W(\xi_i)}} \tag{20}$$

and the error bar on the expectation $A$ is estimated using a Taylor series expansion truncated at the first order term as

$$\sigma_A^2 = \sum_{i=1}^{N_{bulk}} (\frac{\partial A}{\partial W(\xi_i)})^2 \sigma^2_{W(\xi_i)} = \sum_{i=1}^{N_{bulk}} (\beta P_i(A - e^{-\beta u_{B_{1,c}}(\xi_i)})^2 \sigma^2_{W(\xi_i)} \tag{21}$$

and

$$P_i = \frac{e^{-\beta W(\xi_i)} \Delta\xi_i}{\sum_{i=1}^{N_{bulk}} e^{-\beta W(\xi_i)} \Delta\xi_i} \tag{22}$$

where $\sigma_{W(\xi_i)}$ is the error bar on the PMF $W(\xi_i)$ reported by MBAR.

**Table S1.** Overview of MD simulation and analysis method details in this study. We note that for PYL2-ABA binding simulations, the number of atoms and box size vary among parallel trajectories.

| **System** | **Method** | **Software** | **$\#_{atoms}$** | **Size (Å$^3$)** | **Ensemble** | **Simulation time ($\mu$s)** |
|---|---|---|---|---|---|---|
| ABA binding to monomer | adaptive sampling, unbiased MD, MSMs | Amber18 | ~32,000 | ~79*68*73 | NPT | 107 $\mu$s |
| ABA binding to dimer | adaptive sampling, unbiased MD, MSMs | Amber14 | ~62,000 | ~85*85*85 | NPT | 116 $\mu$s |
| *apo-apo* PYL2 association | REUS MD | NAMD | 59,994 | 74*85*117 | NPT | >=8 ns/window |
| *holo-apo* PYL2 association | REUS MD | NAMD | 65,075 | 74*92*117 | NPT | >=8 ns/window |
| *holo-holo* PYL2 association | REUS MD | NAMD | 63,930 | 74*90*117 | NPT | >=8 ns/window |
| PYL2-HAB1 association | REUS MD | NAMD | 82,385 | 79*88*142 | NPT | >=8 ns/window |
| PYL2-ABA-HAB1 association | REUS MD | NAMD | 82,419 | 79*88*142 | NPT | >=8 ns/window |

**Table S2.** Summary of adaptive MD simulations of ABA binding to the monomeric PYL2.

| Round | Parallel simulations | Simulation time (ns) | Aggregate ($\mu$s) |
|---|---|---|---|
| 1 | 50 | 100 | 5 |
| 2 | 50 | 100 | 5 |
| 3 | 50 | 100 | 5 |
| 4 | 50 | 100 | 5 |
| 5 | 50 | 100 | 5 |
| 6 | 50 | 100 | 5 |
| 7 | 50 | 100 | 5 |
| 8 | 50 | 100 | 5 |
| 9 | 116 | 100 | 11.6 |
| 10 | 95 | 100 | 9.5 |
| 11 | 65 | 100 | 6.5 |
| 12 | 100 | 100 | 10 |
| 13 | 100 | 100 | 10 |
| 14 | 100 | 100 | 10 |
| 15 | 100 | 100 | 10 |
| Total simulation time: ~107 $\mu$s | | | |

**Table S3.** Summary of adaptive MD simulations of ABA binding to the dimeric PYL2.

| Round | Parallel simulations | Simulation time (ns) | Aggregate ($\mu$s) |
|---|---|---|---|
| 1 | 50 | 108 | 5.4 |
| 2 | 50 | 96 | 4.8 |
| 3 | 50 | 81 | 4.05 |
| 4 | 44 | 100 | 4.4 |
| 5 | 150 | 108 | 16.2 |
| 6 | 48 | 150 | 7.2 |
| 7 | 86 | 120 | 10 |
| 8 | 93 | 100 | 9.3 |
| 9 | 56 | 100 | 5.6 |
| 10 | 51 | 100 | 5.1 |
| 11 | 111 | 90 | 10 |
| 12 | 200 | 80 | 16 |
| 13 | 100 | 110 | 11 |
| 14 | 100 | 100 | 10 |
| Total simulation time: ~116 $\mu$s | | | |

**Table S4.** Featurization metrics used for the monomeric and the dimeric PYL2 simulation datasets. For the dimeric PYL2, the same set of features were used for both monomers, leading to 64 distances in total.

| Gate loop conformation | | | | | | | | |
|---|---|---|---|---|---|---|---|---|
| | V87:CA | I88:CA | S89:CA | Q90:CA | L91:CA | P92:CA | A93:CA | S94:CA |
| L110:CA | 1 | 2 | 3 | 4 | 5 | 6 | 7 | 8 |
| E147:CA | 9 | 10 | 11 | 12 | 13 | 14 | 15 | 16 |
| Latch loop conformation | | | | | | | | |
| | H119:CA | | H119:NE2 | | R120:CA | | R120:NH2 | |
| L110:CA | 17 | | 18 | | 19 | | 20 | |
| E147:CA | 21 | | 22 | | 23 | | 24 | |
| ABA molecule position | | | | | | | | |
| | ABA: C in -COOH | | | | ABA: O in -CO | | | |
| L110:CA | 25 | | | | 26 | | | |
| E147:CA | 27 | | | | 28 | | | |
| ABA-receptor interaction | | | | | | | | |
| | ABA: C in -COOH | | | | ABA: O in -CO | | | |
| K64:NZ | 29 | | | | 30 | | | |
| Y124:OH | 31 | | | | 32 | | | |

**Table S5.** Reference values of the restrained CVs and the corresponding force constants used in separation PMF calculations for the *apo-apo*, the *holo-apo* and the *holo-holo* PYL2 receptors.

| CV | $k_{force}$ | *apo-apo* | *holo-apo* | *holo-holo* |
|---|---|---|---|---|
| $B_{1,c}$ | 10 kcal·mol$^{-1}$·Å$^{-2}$ | 0 Å | 0 Å | 0 Å |
| $B_{2,c}$ | 10 kcal·mol$^{-1}$·Å$^{-2}$ | 0 Å | 0 Å | 0 Å |
| $B_{1,res}$ | 10 kcal·mol$^{-1}$·Å$^{-2}$ | 0 Å | 0 Å | 0 Å |
| $B_{2,res}$ | 10 kcal·mol$^{-1}$·Å$^{-2}$ | 0 Å | 0 Å | 0 Å |
| $\Theta$ | 0.1 kcal·mol$^{-1}$·$^{\circ-2}$ | 99.391° | 103.461° | 95.735° |
| $\Phi$ | 0.1 kcal·mol$^{-1}$·$^{\circ-2}$ | 101.214° | 100.294° | 110.828° |
| $\psi$ | 0.1 kcal·mol$^{-1}$·$^{\circ-2}$ | -80.633° | -81.441° | -60.528° |
| $\phi$ | 0.1 kcal·mol$^{-1}$·$^{\circ-2}$ | 101.389° | 103.595° | 118.810° |
| $\theta$ | 0.1 kcal·mol$^{-1}$·$^{\circ-2}$ | 96.459° | 101.667° | 94.149° |
| $r$ | 10 kcal·mol$^{-1}$·Å$^{-2}$ | / | / | / |

**Table S6.** Reference values of the restrained CVs and the corresponding force constants used in separation PMF calculations for the PYL2-HAB1 and the PYL2-ABA-HAB1 complexes.

| CV | $k_{force}$ | PYL2-HAB1 | PYL2-ABA-HAB1 |
|---|---|---|---|
| $B_{PYL2,c}$ | 10 kcal·mol$^{-1}$·Å$^{-2}$ | 0.728 Å | 1.508 Å |
| $B_{HAB1,c}$ | 10 kcal·mol$^{-1}$·Å$^{-2}$ | 0.853 Å | 1.812 Å |
| $B_{PYL2,res}$ | 10 kcal·mol$^{-1}$·Å$^{-2}$ | 0.875 Å | 1.204 Å |
| $B_{HAB1,res}$ | 10 kcal·mol$^{-1}$·Å$^{-2}$ | 0.931 Å | 1.812 Å |
| $\Theta$ | 0.1 kcal·mol$^{-1}$·$^{\circ -2}$ | 119.930° | 117.651° |
| $\Phi$ | 0.1 kcal·mol$^{-1}$·$^{\circ -2}$ | 110.142° | 107.921° |
| $\psi$ | 0.1 kcal·mol$^{-1}$·$^{\circ -2}$ | 106.119° | 114.049° |
| $\phi$ | 0.1 kcal·mol$^{-1}$·$^{\circ -2}$ | 76.054° | 77.983° |
| $\theta$ | 0.1 kcal·mol$^{-1}$·$^{\circ -2}$ | 114.946° | 118.954° |
| $r$ | 10 kcal·mol$^{-1}$·Å$^{-2}$ | / | / |

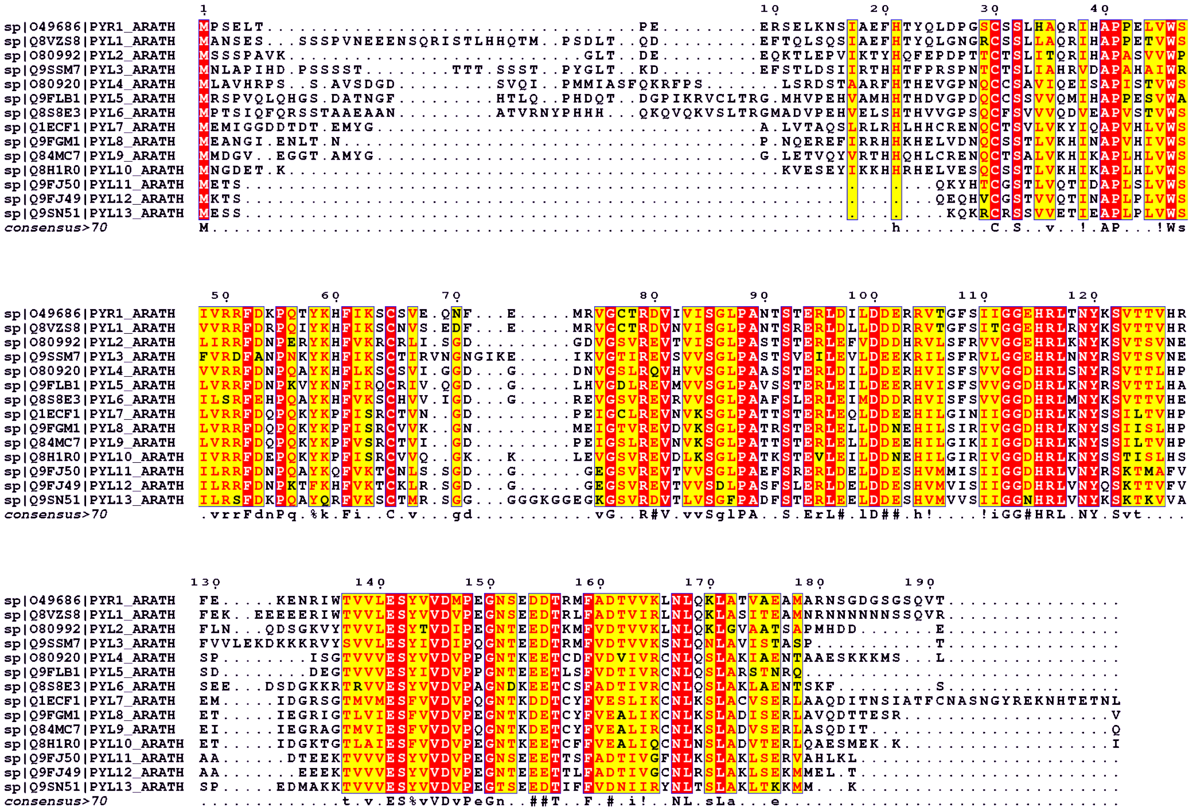


**Fig. S1.** Multiple sequence alignment of the sequences of 14 ABA receptors (PYR1, PYL1-13) in *Arabidopsis thaliana*. PYL1-13 share 50-73% sequence similarity and 38-64% sequence identity as compared to PYR1.

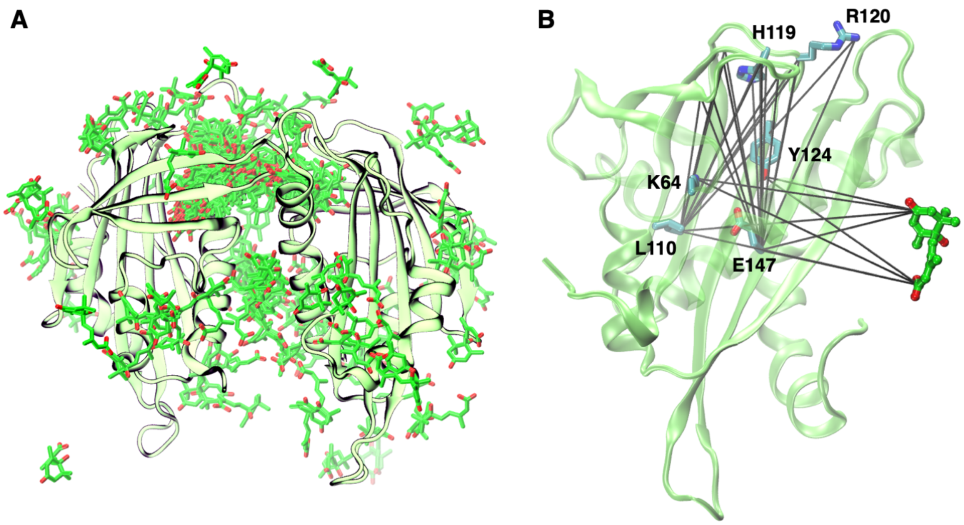


**Fig. S2.** (A) Overlay of ABA positions for all the 200 MSM states of the dimeric PYL2. Our MD simulations have captured the binding of ABA to the left protomer, while also explored the states where ABA resides at top of the binding site in the right protomer. Our analysis was focused on the dynamics of the left protomer. (B) Visualization of the 32 distances used for clustering the PYL2-ABA conformations, which can describe the movement of ABA, the conformational changes in the gate loop and the latch loop and the PYL2-ABA interactions.

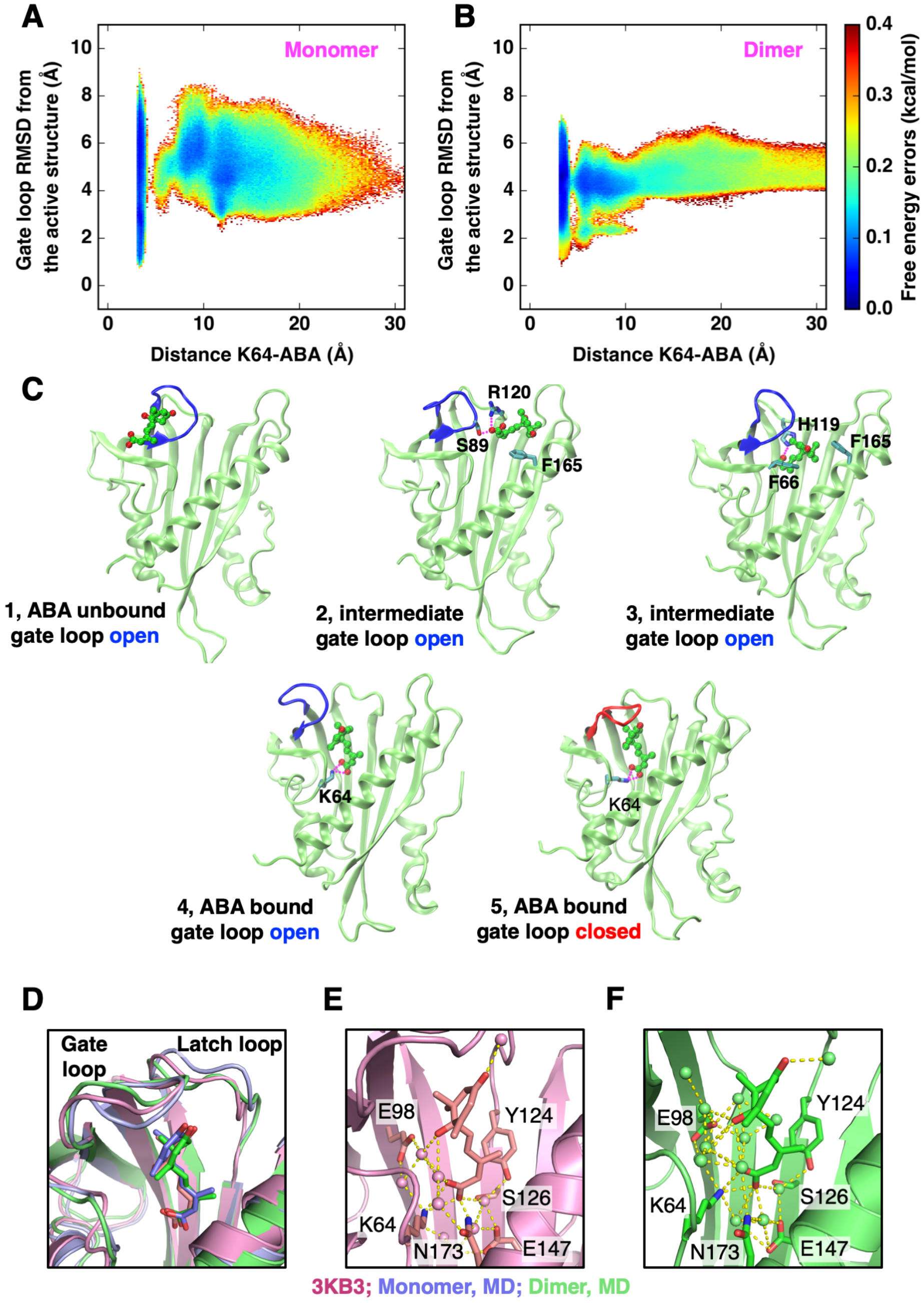


**Fig. S3.** Snapshots of the intermediate states and the active states of the monomeric and the dimeric PYL2 receptors. Errors on the free energy landscapes of (A) the monomeric and (B) the dimeric PYL2 receptors. The errors are the standard deviations of 100 landscapes generated with the projection of random 50% simulation data, weighted by Bayesian MSM probabilities. (C) Structures corresponding to the states on the monomeric PYL2 landscape. The key residues in the PYL2 receptor that mediate the interaction with ABA and the hydrogen bonds formed between PYL2 and ABA are shown. The open (iceblue) and closed (red) gate loop conformations are highlighted. (D) Overlay of the active PYL2 crystal structure (PDB: 3KB3) and the predicted active structures of the monomeric (iceblue) and the dimeric (pink) PYL2 receptors. Water-mediated interactions formed between ABA and the polar residues in PYL2 captured from (E) the crystal structure and (F) the MD predicted structure.

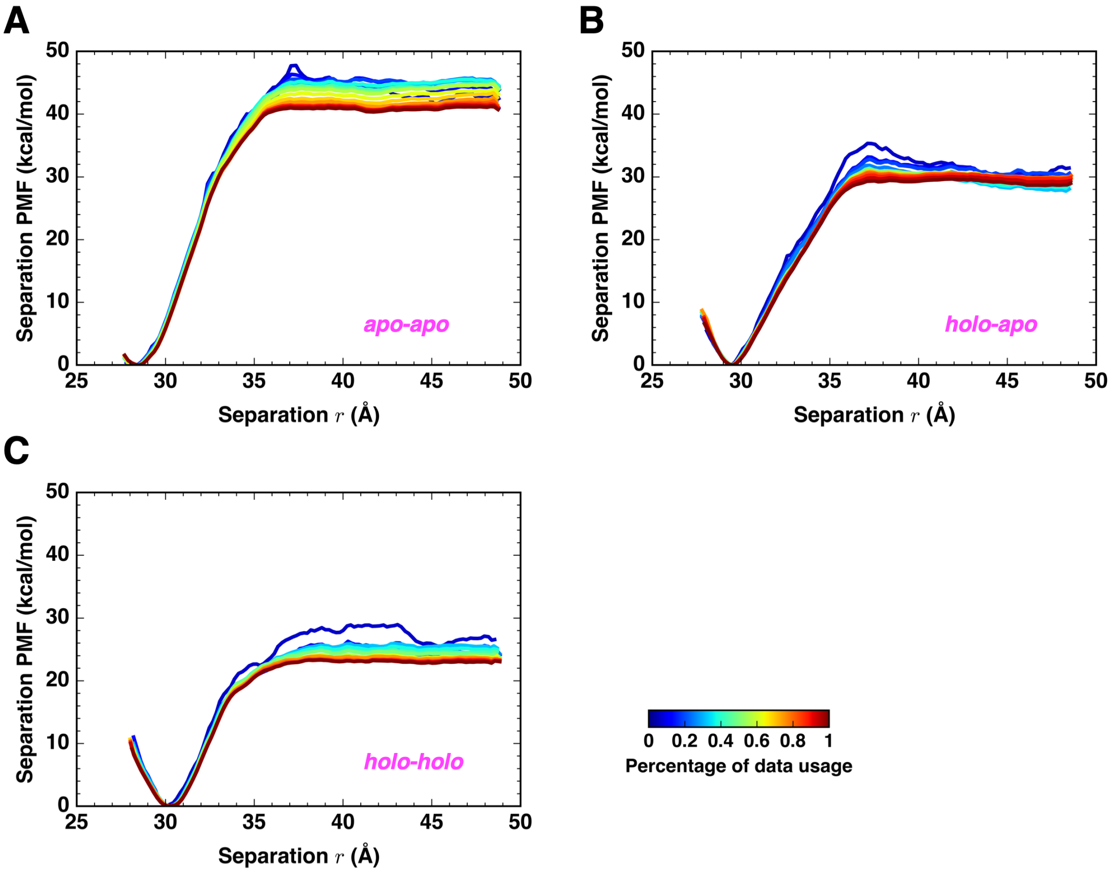


**Fig. S4.** Convergence of the separation PMFs with respect to simulation time. Plots of PMF profiles estimated from the increasing amounts of simulations for (A) the *apo-apo*, (B) the *holo-apo*, and (C) the *holo-holo* PYL2 receptors. The full length of REUS MD simulation for each window is 8 ns.

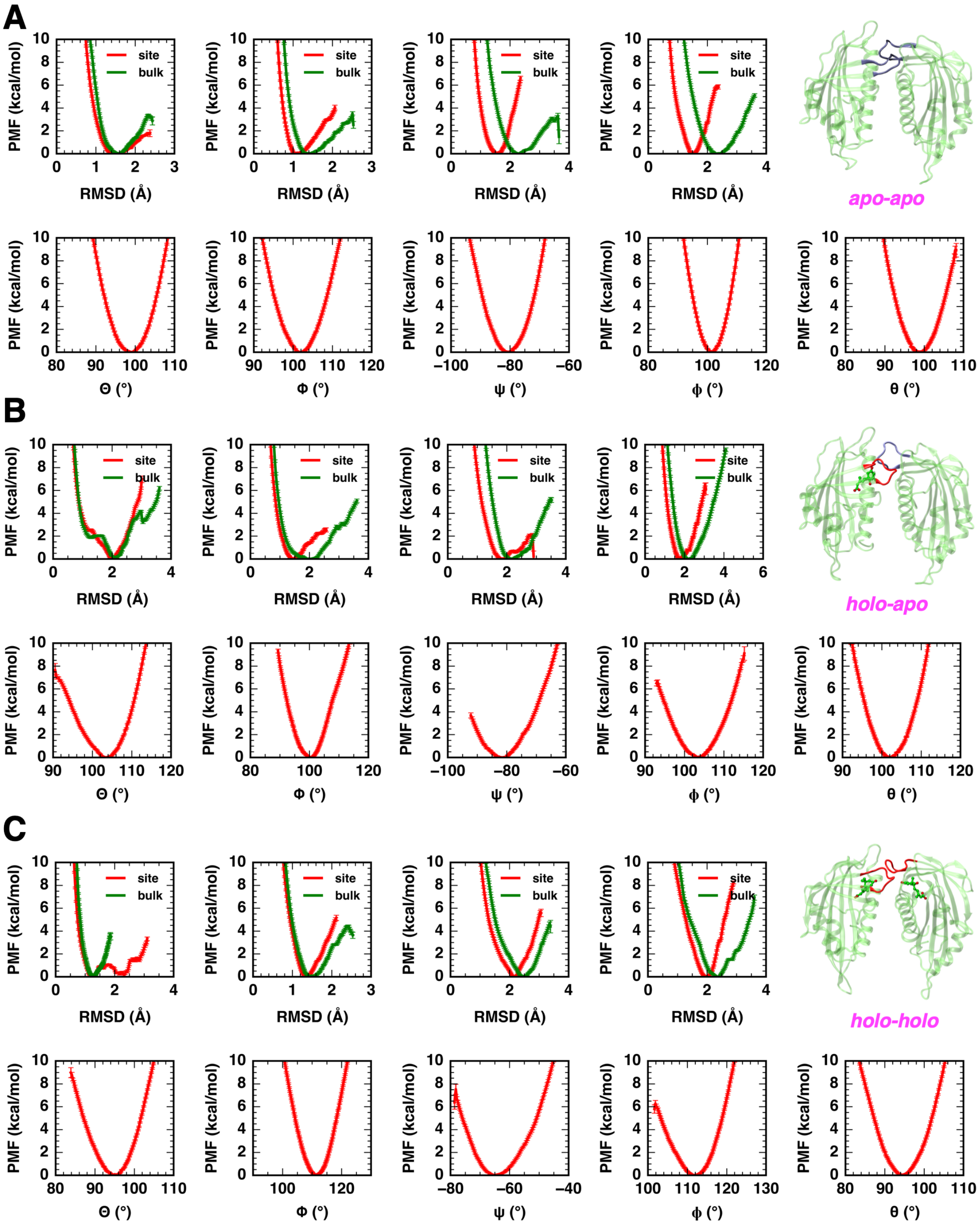


**Fig. S5.** Individual PMFs for all restraint components on (A) the *apo-apo*, (B) the *holo-apo*, and (C) the *holo-holo* PYL2 receptors. Plots for the conformational and orientational restraints $B_{1,c}^{site}$ and $B_{1,c}^{bulk}$, $B_{2,c}^{site}$ and $B_{2,c}^{bulk}$, $B_{1,res}^{site}$ and $B_{1,res}^{bulk}$, $B_{2,res}^{site}$ and $B_{2,res}^{bulk}$, $\Theta$, $\Phi$, $\psi$, $\phi$ and $\theta$ are shown along with the error bars on the PMFs. Snapshots of the starting PYL2 structures are shown.

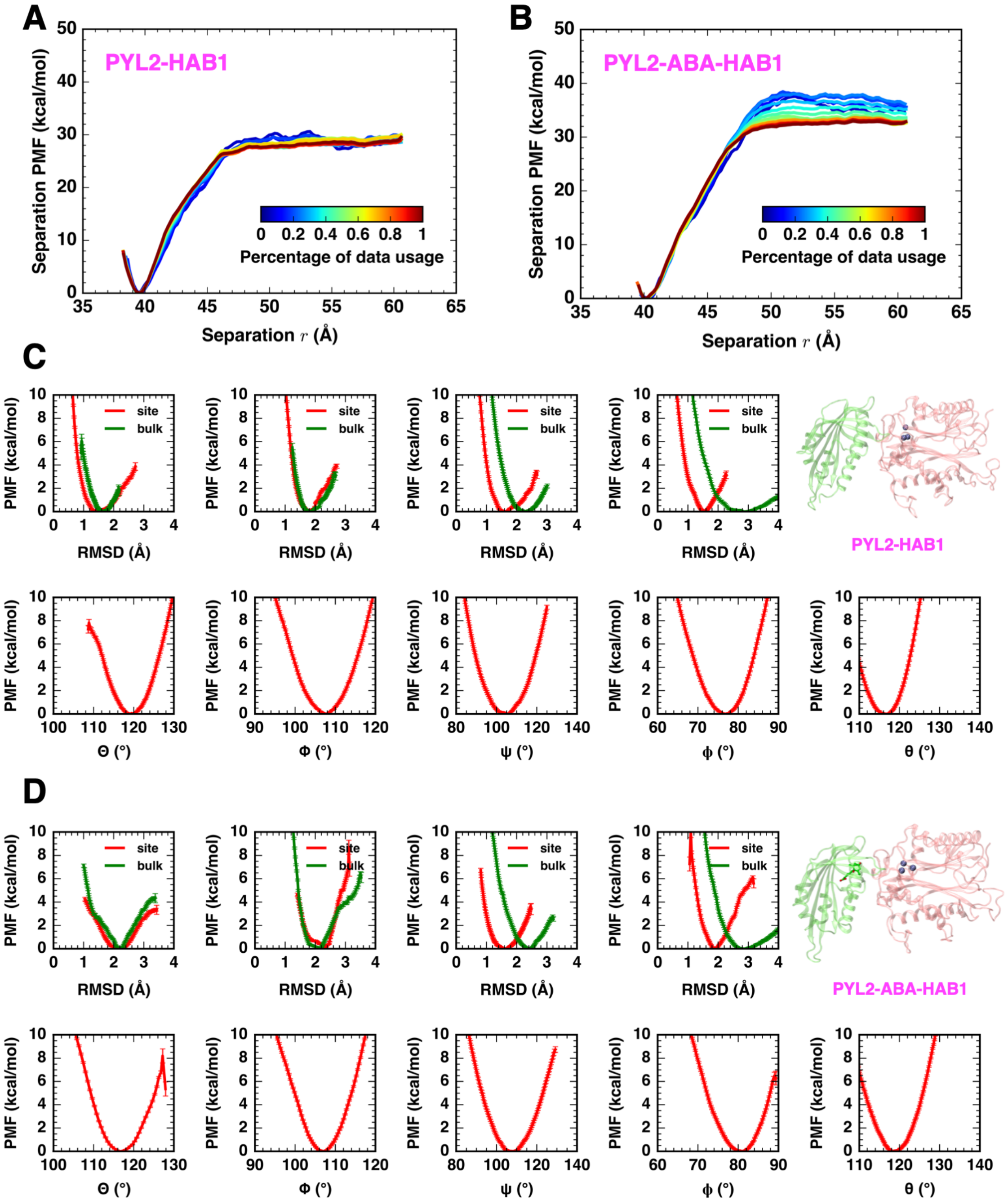


**Fig. S6.** Convergence of the separation PMFs and individual restraint PMFs for the PYL2-HAB1 and PYL2-ABA-HAB1 complexes. Plots of PMF profiles estimated from the increasing amounts of simulations for (A) the PYL2-HAB1 and (B) the PYL2-ABA-HAB1 complexes. The full length of REUS MD simulation for each window is 8 ns. Plots for the conformational and orientational restraints $B^{site}_{PYL2,c}$ and $B^{bulk}_{PYL2,c}$, $B^{site}_{HAB1,c}$ and $B^{bulk}_{HAB1,c}$, $B^{site}_{PYL2,res}$ and $B^{bulk}_{PYL2,res}$, $B^{site}_{HAB1,res}$ and $B^{bulk}_{HAB1,res}$, $\Theta$, $\Phi$, $\psi$, $\phi$ and $\theta$ are shown for (C) the PYL2-HAB1 and (D) the PYL2-ABA-HAB1 complexes. The errors bars on the PMFs are shown. Snapshots of the starting complex structures are shown.

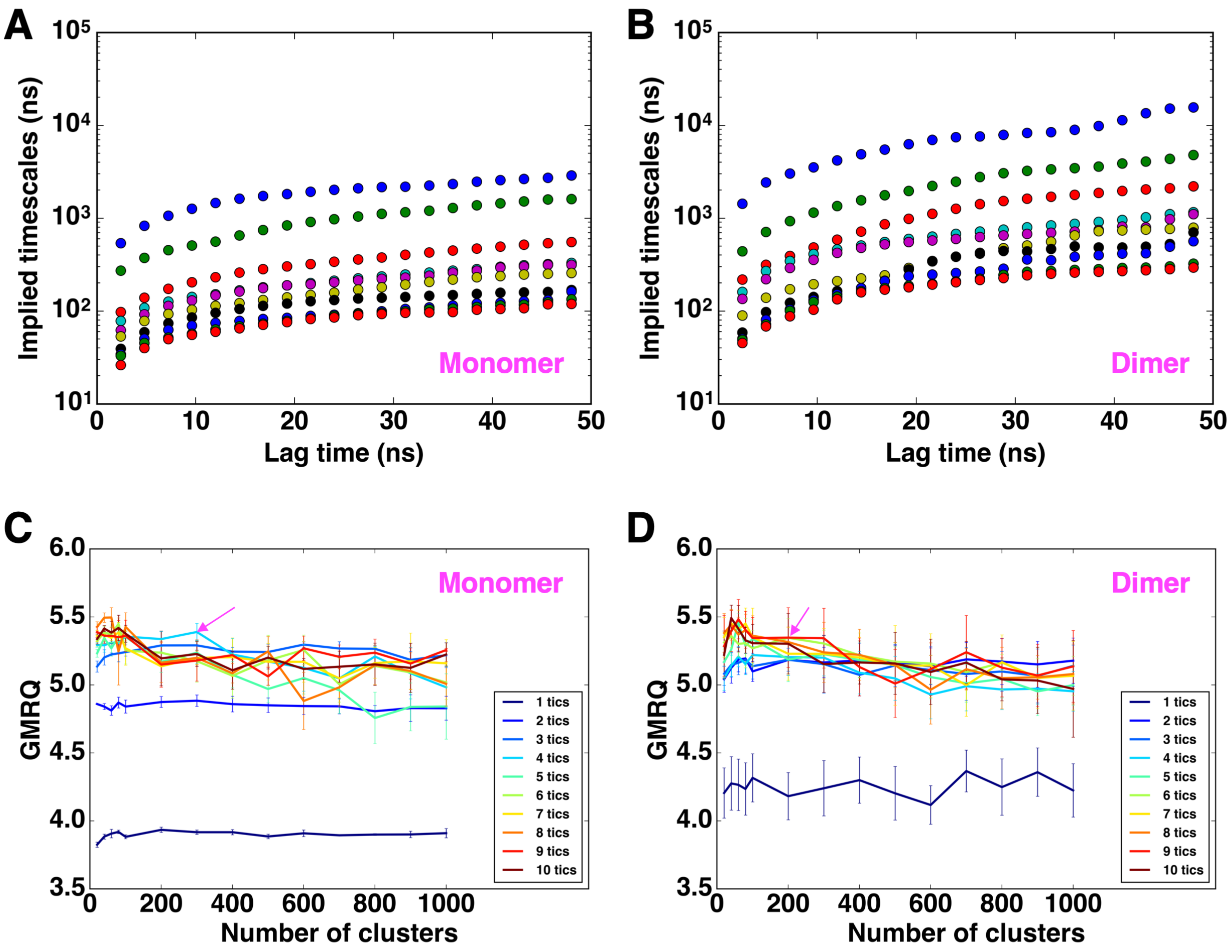


**Fig. S7.** Search of optimal MSM parameters. The implied timescales of the MSMs over (A) the monomeric PYL2 simulation datasets and (B) the dimeric PYL2 simulation datasets. The timescales converge to the true relaxation timescales with the increase of lag time. The lag times used in this study were chosen based on the convergence of implied timescales, which are 40 ns for both systems. GMRQ scores of the MSMs over (C) the monomeric PYL2 simulation datasets and (D) the dimeric PYL2 simulation datasets, with varying number of clusters and tICs. The arrows point to the final MSM parameters used in this study.
